# The structure of full-length AFPK supports the ACP linker in a role that regulates iterative polyketide and fatty acid assembly

**DOI:** 10.1101/2024.07.22.604177

**Authors:** Heidi L Schubert, Feng Li, Christopher P. Hill, Eric W. Schmidt

## Abstract

We present the full structures of two animal fatty acid synthase (FAS)-like polyketide synthases (AFPKs), PKS1 and PKS2 from *Elysia chlorotica.* Unlike the related FAS enzymes that use malonate to produce reduced lipids, EcPKS1 and EcPKS2 accept methylmalonyl-CoA to produce oxidized polypropionate products. When incubated with inhibitory malonyl-CoA (MC), the resulting EcPKS2(MC) structure revealed MC bound to the acyltransferase active site and the phosphopantetheinylated acyl carrier protein (ACP-pPant) bound to the ketosynthase (KS) active site. Remarkably, the entire linker from one modifying region to the ACP was visible in full-length EcPKS2(MC), revealing an asymmetric mega-enzyme structure. Mutations disrupting the affinity between the ACP linker and modifying domains altered substrate selectivity and active site selection, despite an expectation that the interactions are transient and that the wild-type linker alternates between the observed ordered and disordered conformations during the competing catalytic cycles of the homodimeric protein. A second structure, EcPKS2(AC), was acylated both on the KS catalytic cysteine and on the ACP-pPant. The ACP was docked at the dehydratase site, revealing further interactions between ACP and these FAS-like enzymes. The results suggest an unexpected role for the ACP linker in the control of substrate and product selectivity across the AFPK/FAS clade in animals and highlight ACP interfaces and mega-enzyme dynamics over the course of the catalytic cycle.

## INTRODUCTION

Fatty acids and the more complex and diverse polyketides comprise major lipids, clinically relevant therapeutics, photosynthetic protectors, predator poisons, pigments, and many other types of compounds^1–3^. Polyketide synthases (PKSs) and the related cytoplasmic fatty acid synthases (FASs) in animals are mega-enzymes that synthesize these small molecules starting from small acyl-CoA derivatives^1,4–8^. Typically, substrates are selected by an acyltransferase domain (AT) and condensed together in the ketosynthase domain (KS). An associated set of ketoreductase (KR), dehydrogenase (DH), methyltransferase (MT), enoylreductase (ER) and thioesterase (TE) domains potentially modifies intermediates. A 4’-phosphopantetheine (pPant) moiety is post-translationally covalently attached to an acyl carrier protein (ACP), which traditionally has been thought of as a carrier that moves the growing intermediate between catalytic active sites. More recently, attention has focused on potential roles of ACP in protein-protein interactions, such as active site selection and catalytic dynamics, which contribute to processivity and avidity^9–12^. Although it is known that ACPs play a fundamental role in these complex catalytic cycles, the precise nature of these interactions is incompletely resolved in existing mega-enzyme structures.

The diverse family of type I PKS enzymes differ in the number and type of catalytic domains, and the substrate selectivity of each^1,6^. Catalytic domains may be active or residual (structural), and some may be used or skipped during a particular step in the synthetic cycle leading to diverse oxidation patterns along the polyketide chains^4,6,13^. Iterative PKSs (iPKSs) use a single set of enzymatic domains to process all substrates and intermediates until the final product is synthesized, while modular PKSs (mPKS) utilize a separate set of enzymatic domains for each intermediate along a directed assembly production line^7,14^. Mammalian FAS (mFAS) is a specialized type I iPKS that condenses multiple malonyl-CoA (MC) substrates and fully reduces the intermediates in each cycle to produce saturated lipids^15^.

Unlike animal FAS (aFAS) enzymes, iPKSs often produce structurally elaborate polyketides. iPKS and mFAS enzymes use a single ACP that shuttles intermediates between two to five distinct enzyme domains, using evolving substrates at each elongation cycle. It is therefore important to understand how ACP itself can accommodate so many different interactions with high fidelity^8–10^. These interactions and the substrate selectivity of each domain dictate the types of active specialized metabolites that are produced, but the rules are poorly understood, and may not apply across subclades. This leaves a critical gap in the field, in which the many iPKS sequences encoded in fungal and animal genomes are unpredictable and difficult to engineer.

A recently described family of widely distributed aFAS-like PKSs (AFPKs), which are intermediate in character between the aFAS and the traditional type I PKSs, potentially offer a solution to better understand the function of iPKSs^16–18^. In contrast to the related mFAS enzymes, AFPKs accept a more diverse array of acyl substrates, and instead of fully reducing each intermediate they select the use of specific catalytic domains to introduce different oxidation states in a regioselective manner, according to product synthesis rules that which have not yet been fully characterized. Greater than 6,300 AFPK enzymes sequences potentially produce a dazzling array of AFPK enzymes, including EcPKS1 and EcPKS2, from the marine kleptoplast mollusc *Elysia chlorotica*, which are ∼75% sequence similar to EcFAS and ∼80% similar to each other^16,17^. Even so, their products differ in their chain lengths, oxidation patterns, and substrate selection. While FASs use MC, EcPKS1 and EcPKS2 specifically use methylmalonyl-CoA (MMC), creating sidechain methyl groups in the natural products. Unlike mFAS, EcPKSs have a nonfunctional ER domain, leading to polyenes. EcPKS1 and EcPKS2 differ from each other in chain length and in their KR domain action, leading to ketones at different positions on the final products (Fig. 1 and Supplementary Fig. 1). As a result, the enzymes produce complex polyketides thought to help keep stolen chloroplasts alive, allowing the sacoglossan molluscs to live for months by photosynthesis alone^19,20^.

**Fig. 1.**
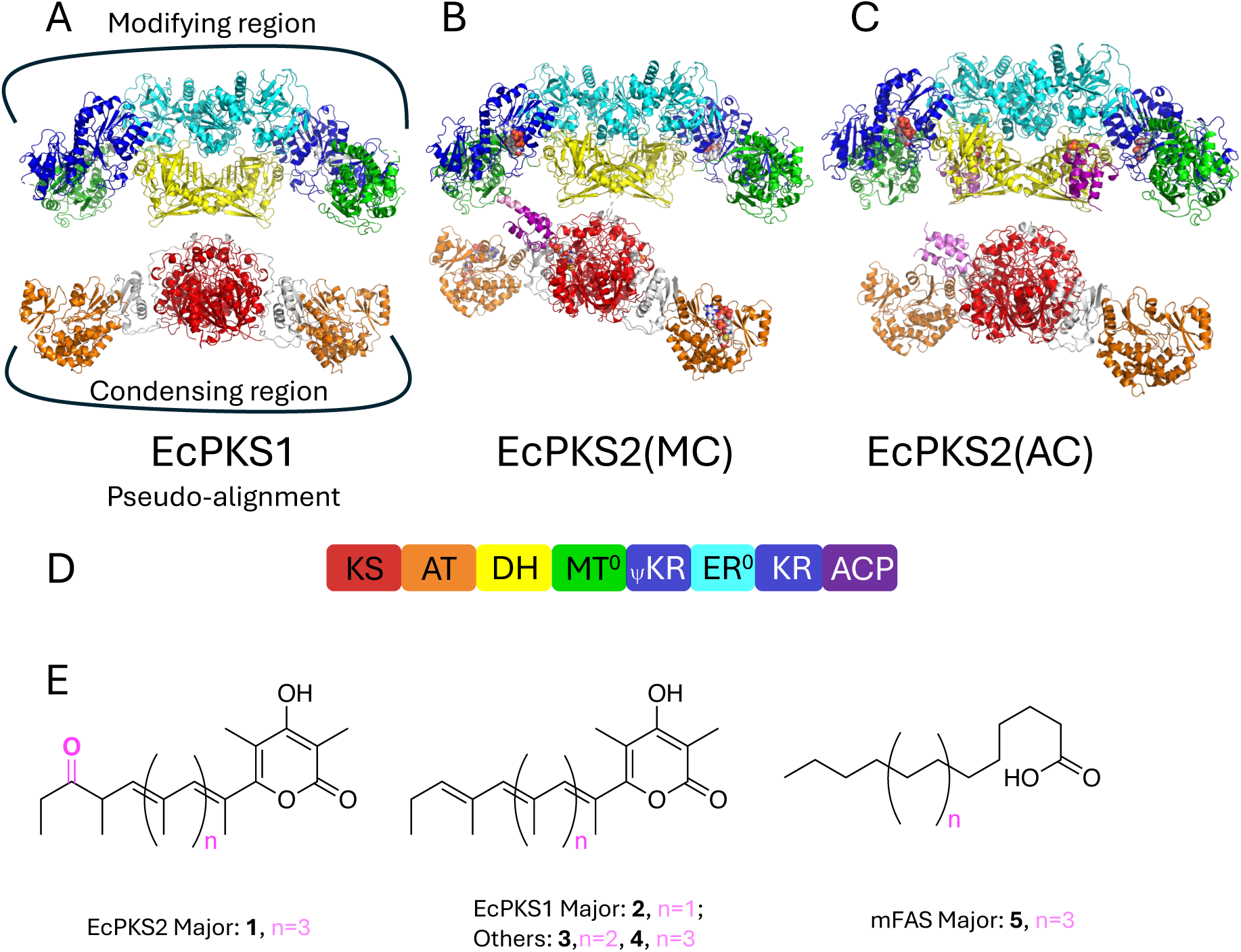
Homodimeric structures and overall schematic of EcPKS1 and EcPKS2. Colors in all structures correspond with schematic in panel D. **A)** EcPKS1 condensing (bottom) and modifying (top) regions in a pseudo-alignment based on EcPKS2(MC). **B)** Structure of full-length EcPKS2(MC). Malonyl-CoA (MC), holo-ACP, and NADPH bound at the AT, KS, and KR active sites, respectively. **C**) Structure of full length EcPKS2(AC). Acylated-Cys187 at the KS domain, ACP-Acetyl-pPant docked at the DH domains and low occupancy ACP at the KS domain. **D**) Schematic of the enzymatic domains. **E**) Primary products of EcPKS1, EcPKS2 and mammalian FAS. Abbreviations: pPant (phosphopantetheine), ketosynthase (KS), acyltransferase (AT), dehydratase (DH), inactive methyltransferase (MT^0^), ketoreductase structural domain (ψKR), inactive enoylreductase (ER^0^), ketoreductase catalytic domain (KR), acyl carrier protein (ACP).

Characterization of the structural framework supporting polyketide production remains challenging^21,22^. Individual AT or KS-AT didomains have been characterized bound to ACP through crosslinking, with synthetic covalent tethers, or as isolated apo-ACP complexes and reveal small charge complementary interfaces^23–30^. Rare full-length structures initially failed to capture the ACP domain due to a flexible linker to the main mega-enzyme, the inherent transient nature of the ACP-pPant for each enzyme interface^31,32^. Recent work on yeast FAS using a large cryoelectron microscopy (cryoEM) dataset parsed out lower occupancy ACP-bound states to reveal several enzyme:ACP interfaces of the unique tandem-ACP within the barrel fungal FAS structure^33,34^. This work highlighted substrate modulation as a tool for capturing the ACP at individual domains, though further work is necessary to explore the co-adaptation of the ACP and enzyme interfaces. In no cases to date has the full linker been visualized connecting the cis-ACP to the mega-enzyme.

The close sequence relationship of the AFPKs with mFAS, coupled with their different product suites (Fig. 1 and Supplementary Fig. 1), provides an unusual opportunity to study how enzyme structural features impact product synthesis. Here, we report the full-length structure determination of both EcPKS1 and EcPKS2 by single particle cryoelectron microscopy (CryoEM). EcPKS2(MC) was determined in the presence of competitive inhibitor MC, which was found bound at the AT catalytic pocket. The inhibition of EcPKS2 at the AT domain is concomitant with the non-covalent capture of the ACP-pPant at the KS domain. A second structure, EcPKS2(AC), was observed with density for a covalently bound acetyl group linked to the active site cysteine of the KS and the ACP-pPant docked at the DH. Remarkably, capture of the ACP at the KS is accompanied by the ordering of the ACP linker as an N-terminal extension of ACP helix α1 resulting in an asymmetric connection between the condensing and modifying regions. Mutation of residues within the modifying region that make contacts with the linker disrupt the catalytic cycle, changing substrate selectivity and inhibition, as well as oxidation patterns in the products. Torsional limitations in the ACP linker itself may impart directionality on the catalytic progression.

## RESULTS

### Structure determination of EcPKS2 and EcPKS1

EcPKS1 and EcPKS2 adopt the canonical “X-shaped” dimeric structure consistent with the close evolutionary relationship between AFPKs and mFAS enzymes (Fig. 1)^16–18,31^. The N-terminal condensing region (KS-AT didomain) tightly dimerizes through a KS/KS’ domain interface, whereas the C-terminal modifying region dimerizes through the DH/DH’, ER^0^/ER^0^’ and DH/ER^0^’ domain interfaces. In all structures the single peptide chain that connects the modifying and condensing domains is poorly resolved and the chain connectivity in the EcPKSs are assumed to be identical to that of the higher resolution mFAS structure^31^. Models and data statistics are in Supplemental Table S1.

For EcPKS1, the structure of the larger modifying region was initially determined using standard single particle CyroEM methods to a final resolution of 3.5Å (C2 symmetry) (Fig. 1a, Supplementary Fig. 2 and 5b). Due to high heterogeneity between individual particles, the N-terminal condensing region was solved only after subtracting the aligned C-terminal modifying region from individual particles. In addition, a low-resolution *ab initio* volume created from the AlphaFold prediction of EcPKS1(17-868) docked into the expected relative position with respect to the modifying region as seen in mFAS, was needed as a starting volume^35^. The resulting 4.3Å reconstruction of the condensing region (C2 symmetry) contained significant differences from the low-resolution starting volume and the AlphaFold model was easily fit into the new density through small rigid-body rotations of multiple sub-domains. The structure of the C-terminal linker-ACP-pPant is not visible.

The EcPKS2(MC) full-length structure in the presence of NADPH and MC was obtained at 3.2Å resolution (C1 symmetry. Fig. 1b and Supplemental Fig. 3, 5c). This showed MC bound in the AT domain and NADPH bound at the KR active site. The ACP-pPant was docked at the KS’ domain active site with a second ACP’:KS’ interaction also visible at low map contour. The linker residues between the modifying domain and helix α1 of the ACP are fully visible in this map but are not visible for ACP’. Further focused, masked and symmetrical C2 refinement of individual condensing and modifying domains improved the resolution to 3.0Å and 2.9Å, respectively (Supplemental Fig. 3 and 5d).

EcPKS2(AC) was determined to final resolutions of 3.0Å and 3.1Å for the condensing and modifying regions respectively after asymmetrical C1 refinement (Supplementary Fig. 4 and 5e,f). Lower resolution full-length maps revealed an asymmetric configuration similar to that of the EcPKS2(MC) structure. Uniquely, on the side of the homodimer with close contacts between condensing and modifying domains, the ACP density is visible at both the KS (pPant not visible) and DH (acetyl-pPant visible) active sites with partial occupancy. On the more open side of the homodimer, ACP’-pPant was observed primarily docked to the DH’ domain. Electron density connected to KS:Cys187 and the pPant sulfhydryl was most consistent with an acetyl group which likely copurified from buffer.

The full-length structure of EcPKS2(MC) is partially rigidified because the condensing region is tethered to the modifying region via the linker-ACP. Nevertheless, CRYOSPAC 3DFlex analysis of EcPKS2(MC) revealed an x-axis wobble of the condensing domain with respect to the modifying domain (Supplemental Movie). Motion is observed radiating out from the fulcrum caused by the ACP-linker tether. No evidence is present for significant y- or z-axis rotations. The motions contained within the unconstrained EcPKS1 dataset are much greater than observed for the EcPKS2 datasets, reminiscent of the earlier mFAS work^11^.

### Condensing region

#### AT domain

The AT domain (EcPKS2 residues 514-833) is separated by a 3β2α-bundle linker domain from the N-terminal KS domain. The AT domain serves as the acceptance site for substrates entering the iPKS cycle and is thus central to understanding how EcPKSs and other AFPKSs prefer methylmalonate rather than the malonate used by aFAS and fungal PKSs. Sequence analyses of bacterial AT domains have shown that residues surrounding the active site serine are predictive of substrate selectivity ^34^. However, these rules do not apply to animal AFPK and PKS sequences, and lack predictive power in animals^17,18,36,37^(Fig. 2c).

**Fig. 2.**
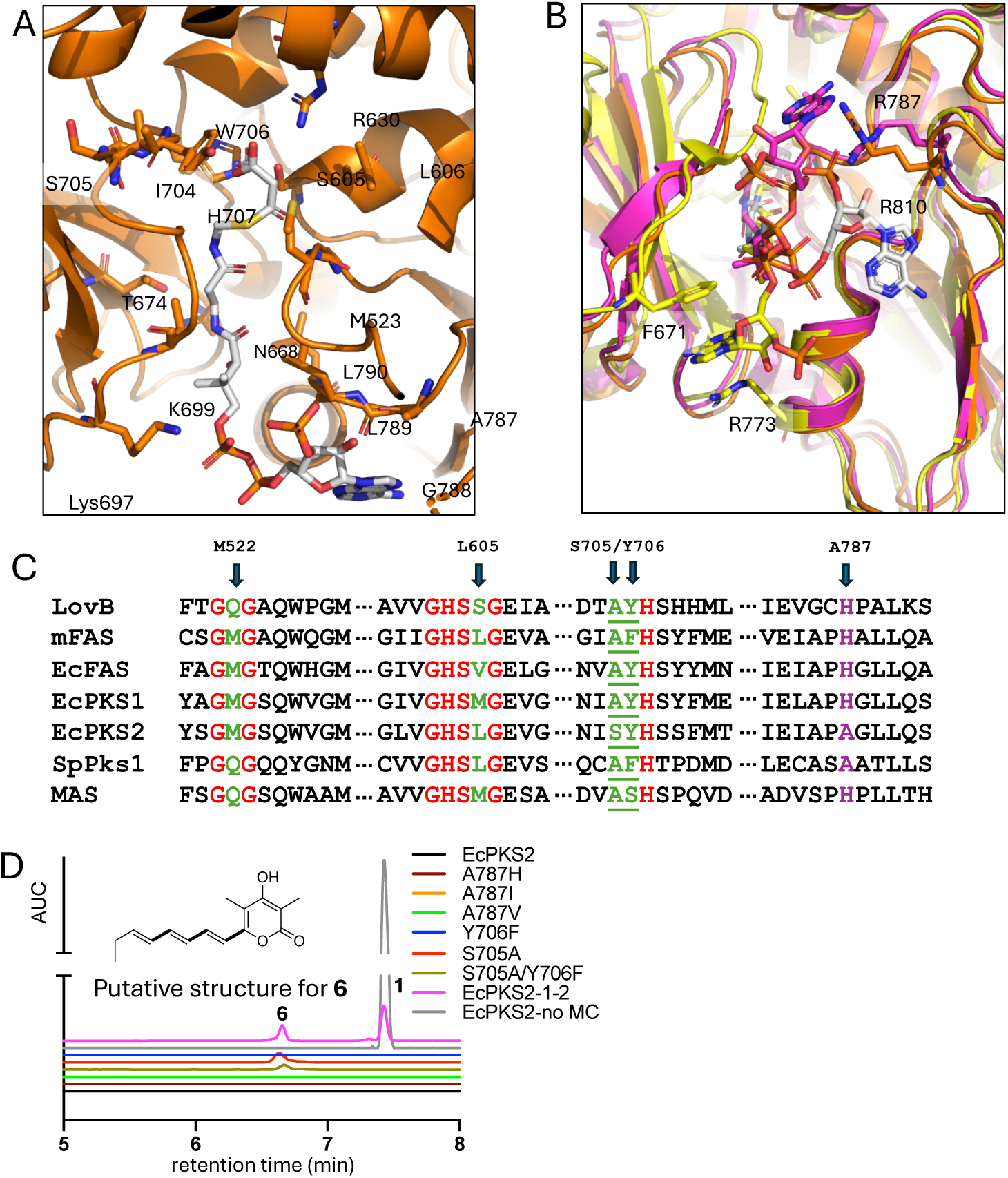
Malonyl-CoA (MC) in the acyltransferase (AT) active site of EcPKS2. **A)** MC (white) bound to the EcPKS2 AT (orange). Active site Ser605 is adjacent to the MC thioester. **B)** Comparison of CoA 3’-phosphoadenosine positions in EcPKS2(MC) (white), murine FAS (PDBcode:6rop, yellow)^25^ and murine FAS (PDBcode:5MY0, magenta)^26^, showing different orientations in all three structures. **C)** Sequence alignment of representative AT sequence motifs thought to have substrate predictive properties in some iPKSs^17,18,36,37^. Arrows indicate regions thought to be important; M522 and L605 are not crucial based upon the substrate selectivity of aligned sequences, and the remaining residues were selected for mutagenesis shown in panel D. **D)** Inhibition of EcPKS2 wild type and mutants after 40 minutes incubation with MC. The graph shows UPLC-MS extracted ion chromatograms (filtered for *m/z* 383.2228 and 245.1183), with the relative area under the curve shown in the y-axis and retention time on the x-axis. In gray, without MC the enzyme produces abundant **1** (product is diluted to 1/50). Production of **1** is completely suppressed by MC in wild type and all AT mutants except for the domain-swapped enzyme EcPKS2-1-2, in which a small amount of **1** is still produced. In some of the mutants, a small amount of product incorporating three units of MC and 3 units of MMC was produced. A possible product **6** is shown; [U-^13^C]-malonate was used to confirm the incorporation of MC (*m/z* 251.1384, Supp. Fig. 9b).

To understand MC inhibition, we incubated EcPKS2 with NADPH and MC. The resulting EcPKS2(MC) structure features MC non-covalently bound to the EcPKS2 AT domain at a local resolution of ∼3.5Å (Fig. 2a and Supplemental Fig 6a). The MC carboxylate moiety interacts with the ring face of active site Tyr706 and experiences charge offset through proximity to His707 and Arg630. Catalytic Ser605 is positioned 3.4Å from the target carbonyl. It is not immediately clear from the EcPKS2(MC) active site why MC is not a major substrate, though the lack of the extra methyl group might twist the substrate slightly, perhaps sufficiently enough to evade nucleophilic attack.

We characterized the affinity of the inhibitor against the wild-type enzyme. EcPKS2 was rapidly inactivated, with a *t*_1/2_ of 1.4 min in the presence of 250 µM MC, with a 10 µM IC_50_ (Supplemental Fig. 7a). Inhibition was competitive with MMC, consistent with the observation of MC in the AT active site in the structure. When glycerol from the enzyme preparation was included in the buffer, inhibition remained competitive but reversible, potentially reflecting greater enzyme mobility.

The large AT active site pocket facilitates only weak interactions (distances between 3.9-4.9Å) with the cysteamine and β-alanine regions of the CoA, including polar interactions with Asn668, Thr674 and Met523. Hydrophobic residues Val699, Leu789 and Leu790 surround the dimethyl carbons of the pantoic acid region, and Lys697 interacts with the first bridge phosphate (Fig. 2a,b). In an mFAS:octanoyl-CoA complex, the nucleotide is packed against the ferredoxin-like domain between residues Phe671 and Arg773, and in the CoA-bound mFAS, where MC has already been transferred to the active site Ser, the nucleotide is at the interface between AT-subdomains packing against conserved Arg787 which plays a dual role in nucleotide positioning and formation of a salt bridge with the bridging phosphate^25,26^. The equivalent arginine in EcPKS2(MC), Arg810, forms a salt bridge with the 3’-phosphate in a very different manner. The 3’-phosphoadenosine group is positioned on the outer surface of the active site pocket interacting with loop residues Ala787-Gly788-Leu789. Critically, the Ala787-Gly788 pair permits close association of the nucleotide with the AT domain unique to other CoA bound structures. To test whether the phosphoadenosine position affects MC inhibition or binding, mutations of EcPKS2 Ala787 to His, Val and Ile were constructed and assayed, but remained inhibited (Fig. 2d).

Residues Ser705-Tyr706 were conserved and corresponded to the V<u>AS</u>H in mycobacterial polyketide synthases that are thought to affect MC selection^39,40^. However, mutation to the more permissive Ala705, Phe706, or Ala705-Phe706 did not alter MC inhibition (Fig. 2d). There was a small but noticeable increase in the incorporation of MC into products, but similar changes were also observed in mutations outside of the AT (see below).

#### KS domain

The KS domain is the site of chain extension and thus plays an important role in determining the chain length of the resulting natural products. Although the EcPKS1 and EcPKS2 KS domains are 83% similar, EcPKS2 produces a major product that is extended by two additional methylmalonate units in comparison to EcPKS1. The EcPKS2(MC) KS domain (residues 16-443 and 845-877) sit with active sites opposed at the central dimer interface of the condensing region (Fig. 1). ACP (residues 2206-2278) is bound at this KS/KS’ interface making interactions through end of ACP(α1), the beginning of ACP(α2), and the middle of ACP(α3). These interactions span the KS:KS’ interface such that ACP(Asp2240:OD1) interacts with KS Gln159:OE1, ACP(Met2243:SC) interacts with KS His94, and ACP(Leu2220:O) interacts with KS Thr95, but ACP(Gln2265 and Leu2242) interact with Asn264’ and Met229’ of the subunit housing the active site residues and a pocket (Fig. 3a). The KS catalytic residues, EcPKS2(His318’, His356’ and Cys187’), are within by the opposing polypeptide chain from the ACP.

**Fig. 3.**
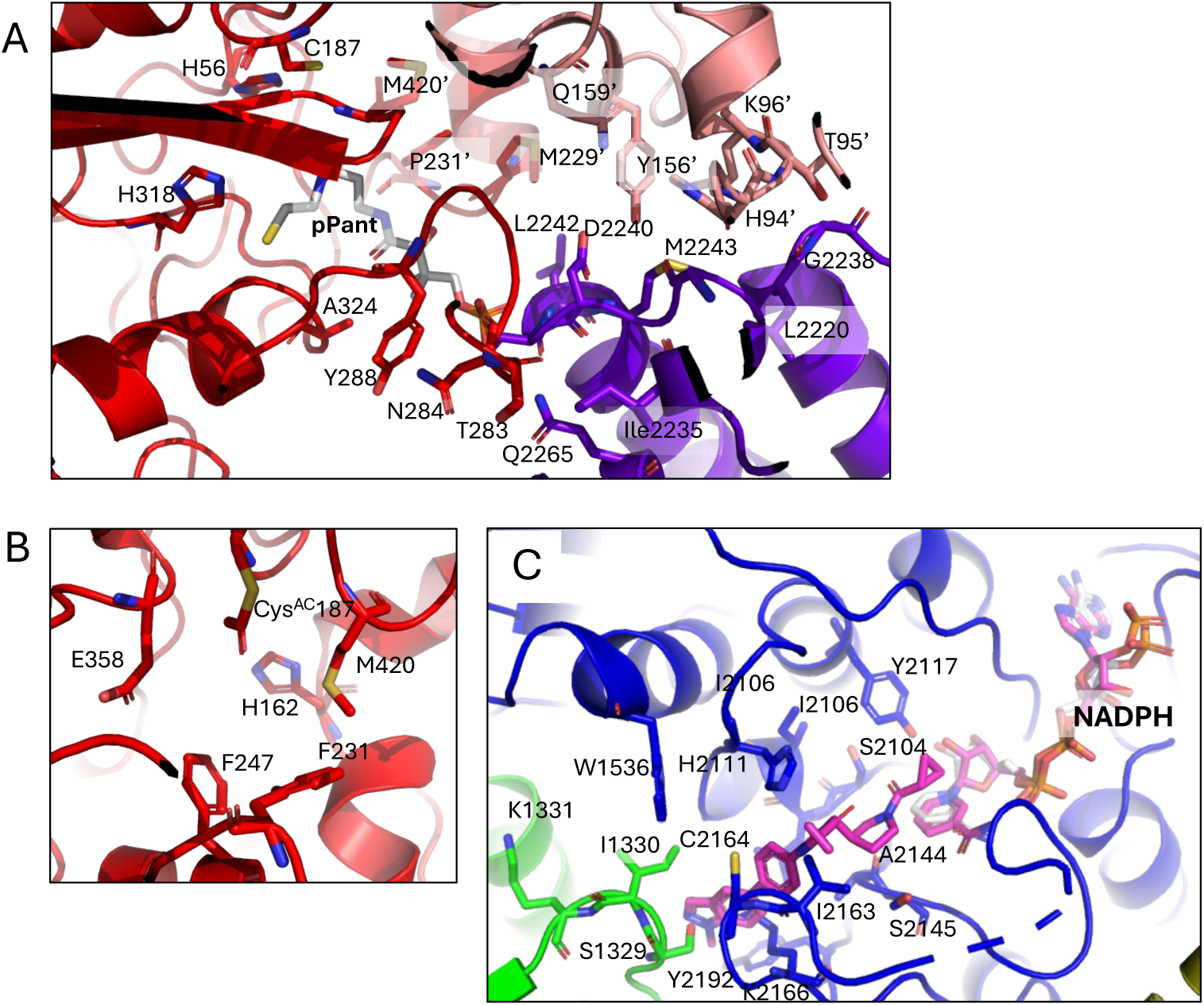
EcPKS2 ketosynthase (KS) and ketoreductase (KR) active sites. (MC) ketosynthase (KS) active site with ACP-pPant bound. **A)** EcPKS2(MC) KS with ACP-pPant bound. Residues surrounding the pPant (white) and at the interface between the KS (red), KS’ (pale red) and ACP (purple). **B)** Residues surrounding the EcPKS2(AC) KS active site acetylated-Cys187. **C**) EcPKS2(AC) ketoreductase (KR) active site (blue). The intermediate binding pocket is highlighted by the location of a spiro-imidazolone inhibitor and NADPH found in hFAS(KR)(PDBcode:5C37, magenta)^27^.

The EcPKS2(MC) covalently bound pPant extends from Ser2241 into the KS pocket towards the Cys187’ active site residue (Fig. 3a). The sulfhydryl group of pPant is in a region of weak density near the active site pocket and is not covalently connected to the enzyme (Supplemental Fig. 6b). Like the malonyl-CoA bound in the AT, no direct hydrogen bonds are formed with the pPant, although a hydrophobic ring comprising EcPKS2 residues Phe231, Tyr288 and Ala324 stabilizes the pantoic acid dimethyl group.

The EcPKS2(AC) structure contains a marked difference at the KS; electron density consistent with the presence of a covalently linked acetyl group is visible on the catalytic cysteine residue (Fig. 3b and Supplemental Fig 6c). This is normally the position in which a starter unit would sit awaiting a translocated extender unit. Only discontinuous electron density is present within the pPant tunnel, indicating that a pPant group is only present in a small fraction of the particles. Nonetheless, a low occupancy ACP is visible outside the KS active site in similar position to that seen in the EcPKS(MC) structure (Fig. 5b,d and Supplemental Fig. 5e).

### Modifying region

#### KR domain

After a short flexible linker from the condensing region, the DH-MT^0^-ER^0^-KR domains dimerize to form the modifying region (Fig. 1). The β-KR domain catalyzes the reaction after Claisen condensation to reduce a β-ketoester to a β-hydroxyester through oxidation of NADPH. In EcPKS2, the first incorporated methylmalonate group retains the ketone, suggesting that the diketide intermediate is not a preferred substrate for KR. Subsequently, four methylmalonate units are added sequentially to the growing chain. At each step, the additional unit is reduced by the KR and dehydrated by the DH to produce a polyolefin. In both EcPKSs, the final three methylmalonate units are not reduced by KR, leading to spontaneous release from the enzyme as pyrones (**1**). In contrast, EcPKS1 reduces the ketone of the first methylmalonate, indicating a difference in substrate selectivity at the KR, or alternatively an intermediate recognition issue at the KS.

Density for NADPH is visible in the EcPKS2 KR active site of both structures, but the flexible substrate-binding loop (2154-2158) is disordered even in the better resolved EcPKS2(AC) structure (Fig. 3c, Supplemental Fig. 6d). EcPKS2(AC) active site residues Tyr2117, Lys2076 and Ser2104 are positioned near the nicotinamide riboside motif, which is positioned similarly to what is observed in the structure of human FAS bound to NADPH and a spiro-imidazolone inhibitor ^41^. In EcPKS2(MC) the NADPH structure sits a bit higher, and the density is much worse for the nicotinamide mononucleotide and substrate-binding loop, emphasizing the differences in conformational flexibility of this region that result from rigidification upon substrate-triggered loop closure – pushing the NADP further into the pocket.

#### DH domain

The DH domain catalyzes a dehydration reaction following the KR domain and eliminates water whenever the KR acts on a specific extender unit, acting similarly in both EcPKS1 and EcPKS2. The DH domain (EcPKS2 residues 896-1132) comprises a pseudo internal repeat of α+β “hot dog” folds. In the EcPKS2(AC) structure, ACP lies below the ER domain pointing the Ser2241:pPant into DH active site, the first time this has been observed for a FAS-like DH domain (Figs. 4,5). Strong density for the pPant is visible past the sulfhydryl group, which lies between the active site residues His903, Asp1058 and Leu1075. An acetyl group has been modeled off of the sulfhydryl, consistent with the density found at the KS active site in this structure (Supplemental Fig. 6e). The pPant pocket is lined with hydrophobic residues including Val905, Ile944, Ala947, Tyr1026, Leu1075, Thr1077, and Ile1127, with the carbonyl oxygens of Ile944 and His945 just beyond (3.5-4Å) hydrogen-bonding distance of the hydroxyl of the pantothenic acid moiety. Mutation of His903 to Ala or Leu1075 to Ser result in shorter pyrone products, which are produced abundantly in previously tested fusion enzymes and apo enzymes^17^ (Supplemental Fig. 10b), indicating that only the condensing region is active.

**Fig. 4.**
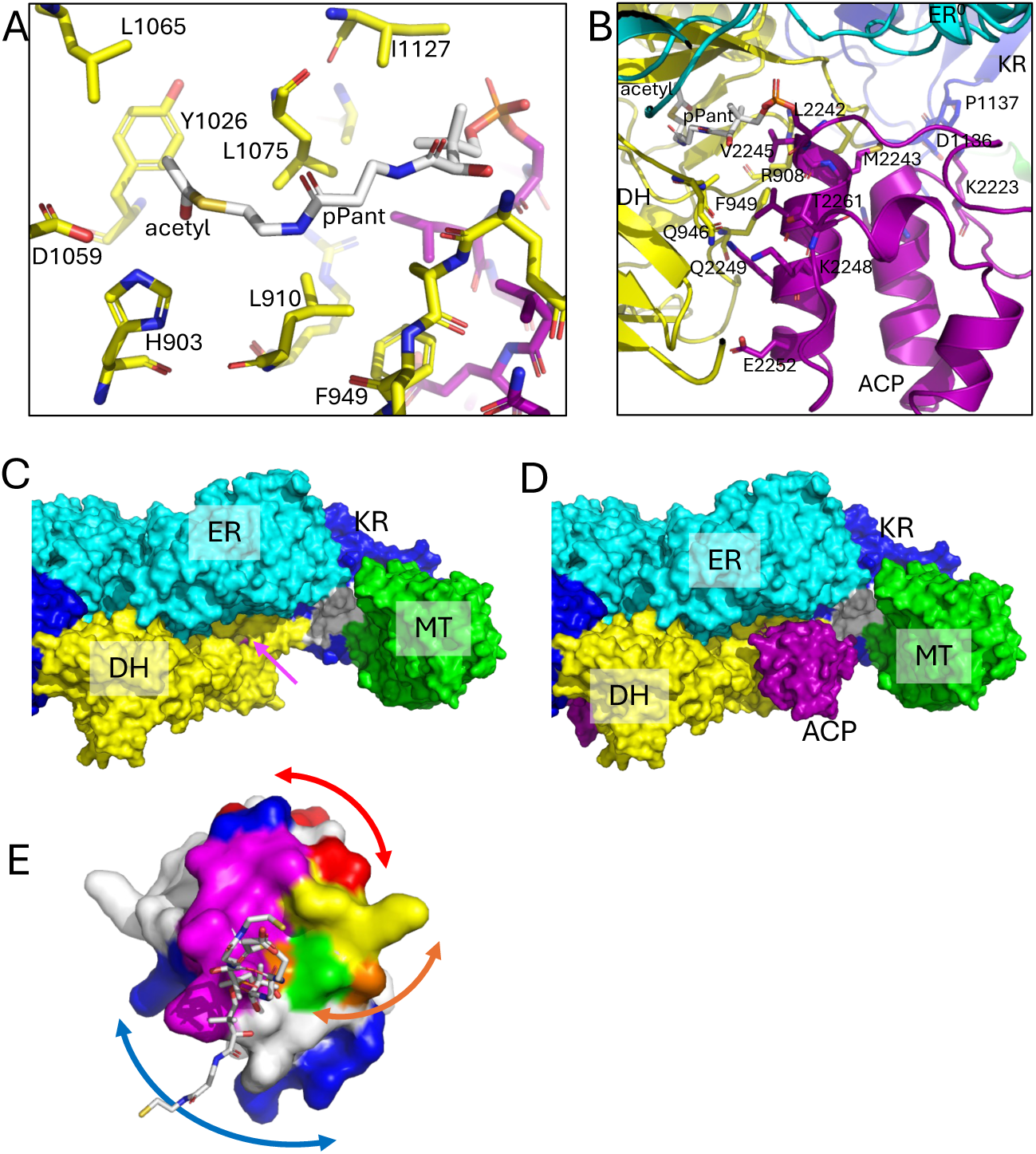
ACP-acetylated-pPant docked at the dehydratase (DH) domain in EcPKS2(AC). **A)** DH (yellow) active site surrounding the acetylated-pPant (white). **B)** ACP (purple) interface with DH (yellow) domain with nearby surfaces of ER^0^ (turquoise) and KR (blue) domains. **C)** Overview of the substrate tunnel in the apo-DH (yellow) domain (pink His903). **D)** overview of the ACP (magenta) docked at the DH (yellow) domain. **E)** Surface of the ACP domain with residues colored by the respective interactions with the AT, KS and DH domains. ACP interactions marked with DH (blue), KS (red) and AT (orange) interactions as well as those residues which overlap at the DH/AT (green), KS/AT (yellow) and DH/KS (green) or all three interfaces (magenta). The pPant shown in white sticks for each docked ACP conformation are shown to highlight the range of conformational diversity.

The ACP:DH interface comprises the surface of ACP(α2) and includes hydrogen bonds from Gln2249(NE2) to Gln946(OE1) and Ala947(O) and GLN2249(OE1) to Phe949(N) as well as van der Waals interactions surrounding the phenyl ring of DH:Phe949 with ACP residues Leu2242, Val2245, Gln2246 and Gln2249 (Fig 4b).

The ACP has now been observed docked to the KS and DH domain within full length FAS/AFPK structures and its conformation can be predicted at the AT domain based on isolated structures (Supplemental Fig. 8a). The three domains make some distinct interactions with a few different residues of ACP but generally overlap in their interaction with the pPant and its nearby region. Charge and hydrophobic surface complementarity dominate the interactions, along with a few hydrogen bonds. The DH:ACP interface resembles the KS:ACP interface described above and has a slightly smaller solvent inaccessible interface of 527Å^2^ while the KS:KS’:ACP interface is 675Å^2^ (Supplemental Fig. 8). Overlapping enzyme residues that bind ACP (Fig. 4e) reveals ACP residues that are specific to binding individual PKS domains.

A 6Å low-pass filter map for the EcPKS2(AC) structure revealed density visible at low contour for the ACP and linker in multiple conformations (Fig. 5b). The linker attached between the KR’ and the DH’-docked ACP’ that is visible at high occupancy extends away from KR’ and joins the start of ACP’:α1 at residue 2207 (Fig. 5c). The same conformation is seen on the other side of the homodimer where both a DH-docked ACP and a KS-docked ACP are visible. The two ACPs are visible at low contour with linkers modeled in a conformation similar to the EcPKS2(AC)DH-ACP and EcPKS2(MC)KS-ACP linkers (Fig. 5d). The EcPKS2(AC) structure highlights the contortions required by the linker to convert from an ordered helical extension to an extended coil. Note that the ACP docked at the KS could not simply swivel down to the DH but would require motions of condensing and modifying regions to provide additional space for that trajectory (Fig. 6).

**Fig. 5.**
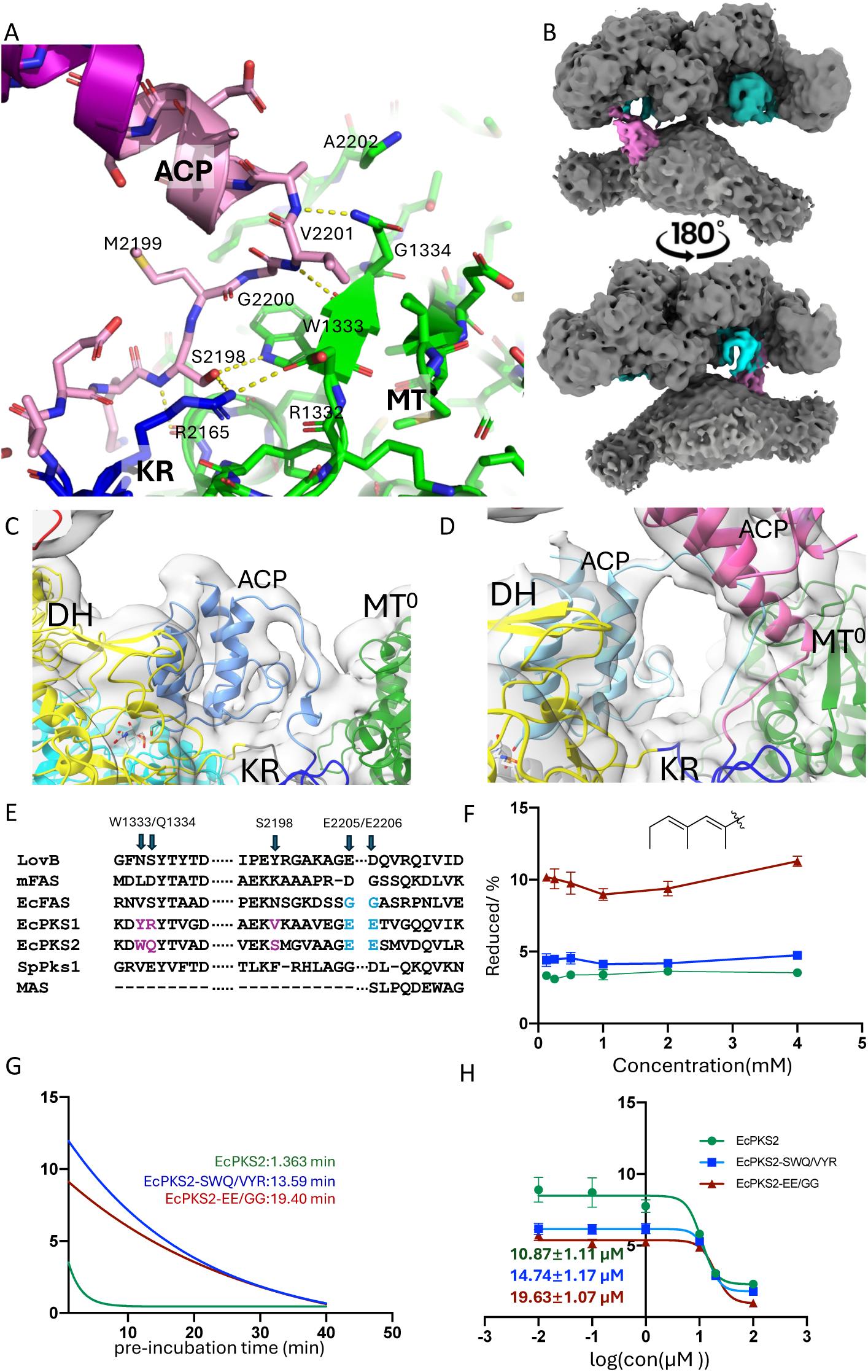
Structure of the ACP linker. **A)** Hydrogen bonds and residues between the EcPKS2(MC) ACP linker and KS and MT^0^ domains. MT^0^ domain (green), KR domain (blue), ACP linker (pink) and ACP domain (purple). **B)** Low contour maps of full-length EcPKS2(AC) revealing three bound ACPs, one at the KS domain like in EcPKS2(MC) (pink) and two docked at the DH domains (shades of cyan). Two panels are flipped 180 degrees on the y-axis to reveal back of protein. **C)** 6Å low-pass filter of the map shown in panel B at low contour to show linker between the KR’ domain and ACP’ docked at the DH’ domain in EcPKS2(AC). **D)** Map in C viewed at the DH, KR, MT^0^ active site revealing partial occupancy ACPs at both the DH and the KS domains with linkers visible to both. **E)** ACP linker (EcPKS2, residues 2198-2206) and converse MT face (EcPKS2 1330-1339) sequence alignment using FAS, iPKS, and AFPK sequences (numbering based upon EcPKS2). These identify linker mutants used in panels F, G and H. **F)** An increase in the percent of reduced products is observed when the ACP linker is mutated. In the y-axis, reduced % was calculated as the ratio of reduced products (**3** and **4**) over the total products, with the concentrations representing AUC measurements made by UPLC-MS. In the x-axis, MMC concentration is shown **G)** Malonyl-CoA (MC) inhibition (*t*_1/2_) of linker mutants with 250 mM MC. The y-axis is area under the curve (AUC) of compounds **1** and **1**’ (structure shown in Supplemental Fig. 7). **H)** MC inhibition of linker mutants. The inset values show the MC IC_50_ in µM for each mutant. The x-axis shows the log[MC] (µM), while the y-axis shows AUC of compounds **1** and **1**’.

**Fig. 6.**
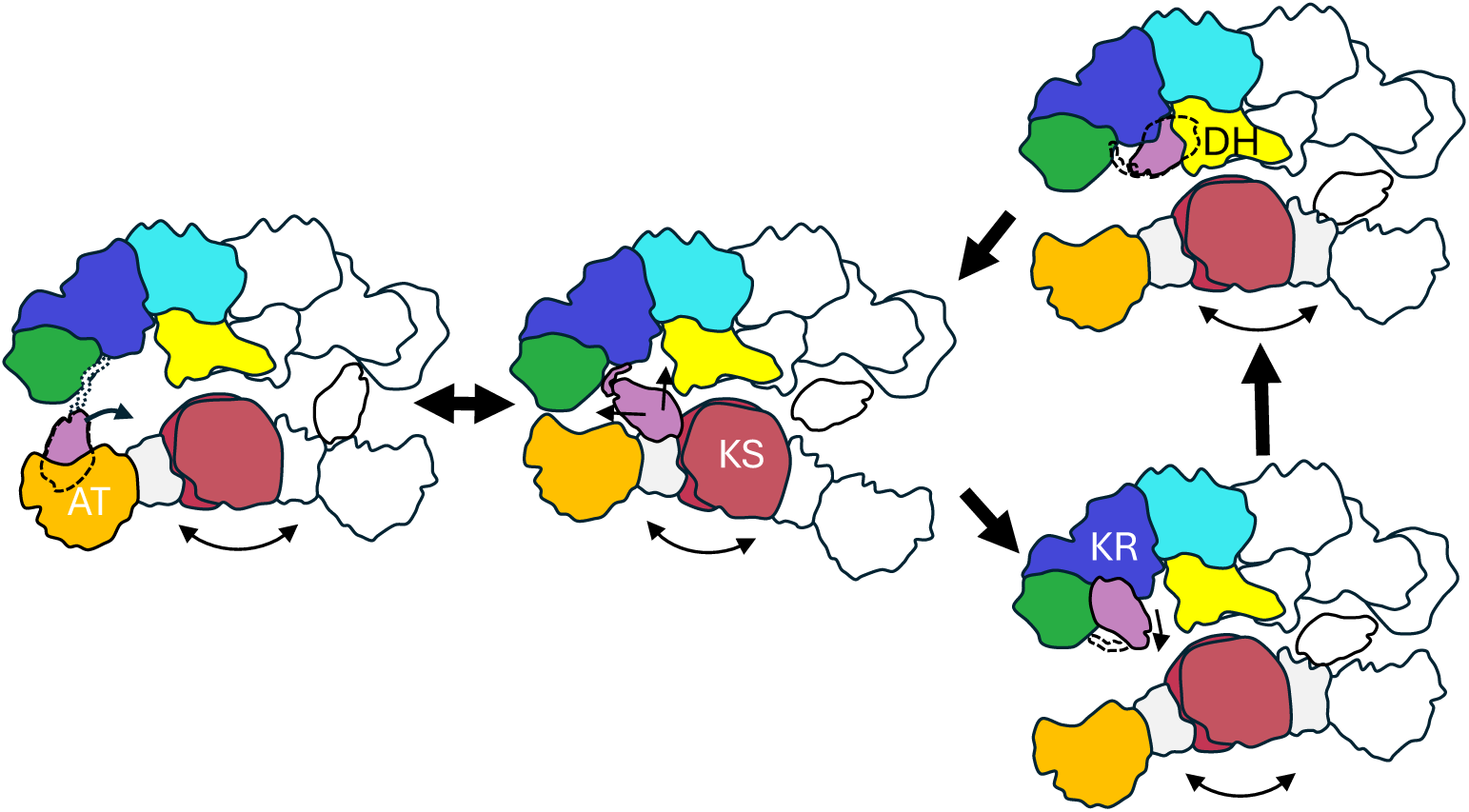
Schematic of domain and ACP movement expected during one side of the mega-enzyme active site cycle. Arrows below the condensing domains reflect necessary motions to allow the ACP to access the front and back sides of the dimer. Dotted lines reflect the flexibility or structure of the ACP-linker and the presence of the ACP at a front-side active site (solid lines) or backside active site (dotted lines).

#### Inactive domains

The MT^0^ and ER^0^ domains are ordered and adopt canonical enzyme conformations but lack active site residues and are not used by the mega-enzymes EcPKS2 and EcPKS1 (Supporting information). Rigid linker between the modifying region and the ACP domain

The ACP domain is increasingly regarded as crucial to substrate processivity. In the FAS/AFPK families, the linker between the KR and ACP domain comprises 8-17 small hydrophilic residues that are thought to be flexible and enable freedom of motion of the ACP. Truncations of this linker and cross-linking have been used as tools to restrict ACP movement and catch ACP docked at active sites^9,28,41^. Many modular PKSs contain additional domains C-terminal to the ACP domain to further restrict and regulate ACP function. Previous reports have not resolved the linker residues, leaving unanswered the question of how the most flexible region of the ACP might participate in producing the diverse array of polyketide and fatty acid derived compounds.

Remarkably, in the full-length EcPKS2(MC), one ACP is rigidly tethered between the KS and MT^0^ structural domain after the end of the KR domain (Fig. 1bc, 5a and Supplemental Fig 6c,e). The ACP linker leaves the KR domain at residue Val2195 and continues in an extended conformation towards the MT^0^ domain to a turn at residue Gly2200 (Fig. 5a). The Ser2198 hydroxyl hydrogen bonds with Trp1333 within the MT^0^ domain and Gly2200 packs tightly against the face of Trp1333. The final linker residues adopt a helical structure, which directly extends into the first helix of the canonical four-helix bundle of the ACP domain. A second ACP at lower occupancy is visibly docked at the opposing KS’ active site, but the linker is not visible (Supplemental Fig. 3) consistent with a mechanism whereby the order and disorder of the linker facilitates the see-saw motion of the condensing region with respect to the modifying region. The AT and KS active sites (and the KR and DH active sites) lie on opposite sides of the central x-y plane and motion of the condensing region is necessary for alternating access to the distant AT active site (Fig. 6). A slight affinity or preference for the KS domain by the ACP, and the ability for the linker to form a ordered structure, may help keep the rotational flexibility in check for enzyme efficiency and substrate progression.

To query the role of the physical interaction between the linker region and the MT^0^ domain, we decreased the interaction interface by mutation. The single residue substitution EcPKS2(Ser2198Val) did not alter substrate kinetics. Slightly larger but still modest effects were seen with two more extensive mutations; Ser2198Val/Trp1333Tyr/Gln1334Arg (EcPKS2-SWQ/VYR) replaced EcPKS2 residues with the corresponding EcPKS1 sequence, and Glu2204Gly/Glu2205Gly (EcPKS2-EE/GG) was intended to imitate the sequence found in EcFAS, which we also hypothesized would disrupt the linker region-MT interactions and increase motility of the ACP (Fig. 5a,e). In the presence of MMC, the k_cat_/K_m_ was slightly altered between the EcPKS2-SWQ/YVR, EcPKS2-EE/GG and the wild type, although the V_max_ was reduced from 51 in wild type to 37 and 23 AUC min^-1^ in the mutants, respectively (Supplemental Fig. 7b). In addition, the product spectrum was similar with all three enzymes, such that compound **1** was the primary product of all three enzyme variants (Supplemental Fig. 10a).

A major difference between these EcPKS2 enzyme variants was that the use of the KR domain was altered. In wild-type assays, the vast majority of product was **1**, which contains a ketone in the first position, and where <4% of products were reduced to the olefin, producing compounds such as **3** and **4**. With EcPKS2-SWQ/YVR, the ratio was only slightly increased, but with EcPKS2-EE/GG a full 10% of products were reduced, representing an easily observed 2-fold increase (Fig. 5f, Supplemental Fig. 7c). A second difference was observed in response to the presence of MC. All three enzymes were inhibited, but the mutants were inhibited to a lesser degree, and incorporated MC into product (**1’**). EcPKS2, EcPKS2-SWQ/YVR, and EcPKS2-EE/GG had IC_50_s of 10, 15, and 20 µM, and *t*_1/2_s at 250 mM of 1.4, 14, and 19.4 min, respectively (Fig. 5g,h). Collectively, these results suggest that the ACP is much more than a carrier of intermediates. When the flexibility of the linker is increased, changes in enzyme active site activity are observed, leading to differences in both products synthesized and inhibitor kinetics. One hypothesis is that the increased flexibility leads to less time at the KS active site and more time sampling other active sites, which leads to spurious KR reduction. Another by-product of dwell-time at the KS active site is that the ketone is left on the wild-type enzyme products. EcPKS2 product **1** is the result of rapid return of the ketone-containing intermediate to the KS active site by the ACP. Once the linker rigidity has been compromised, the ACP might increase the interaction time with other active sites before returning to the KS.

Although the EcPKS1 structure did not display density for the ACP or linker, we made reciprocal mutations in the putative linker-MT regions to more closely imitate those found in EcPKS2: EcPKS1-VYR/SWQ, consisting of Tyr1333Trp, Arg1334Gln, Val2185Ser, and Lys2186Met; and EcPKS1-polySW, consisting of Lys2184Ser, Val2185Ser, Lys2186Ser, Ala2187Ser, Asp1332Trp, Tyr1333Trp, Arg1334Trp, and Tyr1334Trp. EcPKS1-polySW was inactive, but EcPKS1-VYR/SWQ had double the k_cat_/K_m_ in comparison to wild type enzyme (Supplemental Fig. 7b2, 7c). This result reinforced the central importance of the linker region in the overall catalytic cycle of the mega-enzyme.

In previous work, we hybridized EcPKS1 condensing domains with the modifying domains and ACP from EcPKS2, and vice versa. These results strongly implicated the KR as dictating the reduction pattern, and the KS as dictating the chain length^17^. Our new results revealed that the ACP linker region might also play a major role in product formation since more reduced products were obtained when the EcPKS2 linker was mutated. We therefore created a hybrid in which the ACP was also swapped, joined at Val2194 just upstream of the linker region, creating EcPKS2(KS-AT)-EcPKS1(MT^0^-DH-ER^0^-KR)-EcPKS2(ACP), named EcPKS2-1-2. When MMC was used with EcPKS2-1-2, the expected results were achieved: we obtained both the reduced product expected from the EcPKS1 KR, but also the ketone-containing product as expected from the EcPKS2 KR (Supplemental Fig. 9). When MC was added, we saw an increase in malonate-derived products **6** and **2’** (Fig. 2d, Supplemental Fig. 9b, **2’** structures are shown in figure 7c). While some of these were observed in trace amounts from the wild type enzymes, they became dominant products in the mutants. Thus, the overall kinetics of the mega-enzyme complex, governed by the flexibility or the rigidity of the ACP, modifies the substrate and product profile ways that are currently unpredictable.

## Discussion

The discovery of AFPKs bridged an evolutionary gap between FAS and PKS enzymes^16^, and provided model systems for the exploration of substrate selection and intermediate processing experiments relative to both animal FAS and iPKS proteins. Indeed, the global EcPKS2 structure has many similarities with the previously described mFAS, but nonetheless the mega-enzymes make quite different products. In contrast to our initial expectations, differences in active sites could not fully explain the chemistry of the EcPKS enzymes. Instead, our data implicate ACP mobility as a key factor shaping both substrates and products of the AFPK and FAS enzymes.

Previous studies have addressed ACP and PKS motion but have left gaps in understanding. Brignole et al. used negative stain electron microscopy with the porcine FAS to described at low resolution a continuum of orientations suggesting how the ACP of one monomer could access all active sites by rotation of the condensing region with respect to the modifying region^11^. Subsequently, catalytic domains were often studied independently, spliced out of the larger structures for smaller substrate-bound complexes, covalent targeted inhibitors, and crosslinked ACP interactions^23–30^. Smaller mPKS structures like PikAIII shed light into the ACP dynamics, and the complete LovBC complex revealed intradomain interactions of iPKSs^6,42^. Recently data on the barrel structure of yeast FAS was dissected to reveal the unique double-ACP domain docked at each enzymatic active site from large datasets incubated with different ligands to promote stalling the ACP at different locations^33,34^. Finally, iPKS dimer fragility led to Fab-stabilized structures, such as DEBS and Lsd14, which nonetheless revealed the remarkable agility to impede access of the ACP to a KS domain in a turnstile mechanism and consequently funnel substrates forward through the mPKS reaction pathway^42,43^.

The major difference between the observed structures of EcPKS2 in comparison to previously reported structures of mFAS and other iterative iPKS enzymes is the well-structured linker between the EcPKS2(MC) modifying region and the ACP domain that leads to an asymmetrical dimer. This result suggests that linkers may not always be fully disordered. Instead, linkers have potential structure and could preemptively influence the position or rotational direction of the cycling ACP as well as the angle between condensing and modifying domains. In addition, the presence of several structures with different ACP docking poses enabled the definition of binding interfaces, which had previously not been observed in mFAS and type I PKS structures. An underrated part of the ACP surface interface is the pPant itself and closely neighboring ACP residues, which have very little specificity to individual enzyme tunnels yet must add to overall affinity. For example, in EcPKS2(AC)ACP, Gln2249 makes hydrogen bonds with the DH domain. This flexibility supports reproducible placement of the pPant such that longer and longer substrate intermediates are nonetheless inserted deeply into the active sites for catalytic manipulation near the end of the pPant at the thioester-intermediate bond. Collectively, these linker and ACP interactions impact mega-enzyme dynamics and catalysis.

The mechanism of ACP mobility and specificity has been hypothesized as free-diffusion or conformational constraint (medium or excessive)^21,22,44^. In the medium conformational constraint model, some ACP:active site positions are predisposed or preferred through selection mechanisms that are not fully understood, including charge and surface complementation between the ACP and active sites characterized in structural complexes^45^.

The observation of an ordered linker at the KS active site suggests an additional mechanism directing the mobility of ACP, since this particular ACP position maybe occupied more frequently than the other three enzymes. Indeed, this position of ACP is seen in both AT-inhibited EcPKS2(MC) and starter-unit cycled EcPKS2(AC) structures, though an enzyme with just a starter unit might also have affinity at the DH domain.

As the ACPs carry intermediates between the opposing sides of the homodimer, the relationship between the modifying and condensing regions must see-saw back and forth (z-axis) to allow the ACP to pass (Fig. 6). Additional rotational motions along the y-axis appear necessary for the access of ACP to both faces of each domain (in front and behind the image plane drawn in Fig. 1.) The requirement for this motion was directly observed in the transition between the KS-docked pose of EcPKS2(MC) with the DH-docked pose of EcPKS2(AC). The full-length EcPKS2(AC) structure displays asymmetry with the compressed side exhibiting a mixture of KS- and DH-docked ACP and the open side displaying primarily a DH-docked ACP. The structure exemplifies the dynamic and dependent nature of the opposing sides of the homodimer, in which space is needed to cross between active sites that lie on either side of the x-y plane. It is unlikely that the ACP tethered to the KS domain could detach and reach the nearby AT domain without concomitant z-axis rotation of the two regions. Rotational dynamics would permit ACP access from one face of the homodimer (Backside of Fig. 1 for KR and AT active site) to the other (Front side of Fig. 1 for DH and KS active site). During the rotational acrobatics, the linker region would be expected to fold and unfold to permit access to the four active sites, but additional rotational limitations may be imposed due to torsion restraints.

Despite groundbreaking advances, it has remained exceptionally challenging to engineer iterative PKS and FAS enzymes via synthetic biology, or to predict potential structures from sequences, leaving a significant gap in the field. By observing the ACP linker interactions at the molecular level, we reveal an unexpected factor that impacts substrate and product selectivity independent of the preferences of enzyme active sites, though understanding the coevolution of these interfaces could yield an engineered ACP able to adapt to diverse surfaces. While complicating simple models in which enzyme active site residues dictate substrates and products, these results provide avenues for productive discovery and engineering of lipids and polyketides underlying animal and human health.

## Data Availability

Primary CryoEM micrographs and model coordinates will be deposited at the EMDB and PDB databases.

## Acknowledgement

This work was funded by NSF 2203613. We would like to acknowledge use of the University of Utah Electron Microscopy core facility. The support and resources from the Center for High Performance Computing at the University of Utah are gratefully acknowledged. A portion of this research was supported by NIH grant U24GM129547 and performed at the PNCC at OHSU and accessed through EMSL (grid.436923.9), a DOE Office of Science User Facility sponsored by the Office of Biological and Environmental Research.

## Supplemental Movie

3D-flexibility observed within the EcPKS2(MC) dataset. Two latent coordinate series showing motions radiating out from the ACP-linker fulcrum are depicted here. One (in yellow) reveals more y-axis motion compressing and releasing the distant AT domain versus the distant KR/MT while the second (in blue) reveals more x-axis motion rolling the condensing region side to side with respect to the modifying region.

