## Supplemental Information: Methods and Figures for "The structure of full-length AFPK supports the ACP linker in a role that regulates iterative polyketide and fatty acid assembly"

### 1 Experimental section:

#### 1.1 Expression and purification of mutant enzymes

Mutant EcPKS2 and EcPKS1 plasmids were constructed with PCR and yeast recombination using primers (Table S3) in the previously described wild-type EcPKS1 and EcPKS2 expression plasmids (16), followed by verification using total plasmid sequencing (plasmidsaurus). Proteins were expressed and purified using the previously described method (16).

#### 1.2 Cryo-EM specimen preparation, data acquisition, model building and refinement

##### 1.2.1 EcPKS1 structure determination

EcPKS1 at 6.6 mg $ml^{-1}$ was mixed with Tween 20 (1%) to final concentrations of 0.08%. Samples were blotted on AuUltrafoil R1.2/1.3 grids 300 mesh grids after 25 s, 25 mA glow discharge. Grids were blotted for 1-2 s on Mark V vitrobot or 5-6 s on a Leica EM GP2 at 85% humidity and 12 $^{o}$C and plunged into liquid ethane cooled by liquid nitrogen. Data collection occurred on a Titan KRIOS outfitted with a Gatan K3 camera at the University of Utah CryoEM Core facility. Forty frame movies were collected at nominal 81,000x magnification with a pixel size of 1.06 Å/px. Movies received 40-50 e per movie recorded by EPU with a defocus range between -0.08 and -1.8 μM. Data was processed in CRYOSPARC, including 5x5 patch motion correction and CTF analysis. Particles were picked using a 150Å blob size and extracted with a 320Å box initially reduced to 156Å box for preliminary analysis.

EcPKS1 particles were screened by repeated rounds of 2D classification down to 143K particles (Supplementary Fig. 2). De novo 3D volume creation followed by C1 and then C2 refinement led to a high-resolution map dominated by the larger modifying region (3.5Å). Attempts at adding focus masks at the expected position of the condensing region did not provide additional resolution for this region. Subtraction of the masked modifying region from the original particles provided the basis for further reconstruction of the condensing region. De novo 3D volume creation still failed on these maps and a map created from a ColabFold(35,46) prediction of the EcPKS1 condensing region cut to 20Å was also necessary for the successful reconstruction. After masking and application of C2 symmetry the final reconstruction of the condensing region is 4.5Å (Supplementary Fig. 2 and 5). The ColabFold model was docked into the density and rigid body rotations of individual domains optimized the fit. Final real-space refinements were minimal (Supplemental Table S1).

##### 1.2.2 EcPKS2(MC) structure determination

EcPKS2 at 6.8 mg $ml^{-1}$ was mixed with Tween 20 (1%) to a final concentrations of 0.08%. NADPH and malonyl-CoA were added to final concentrations of 1 mM and incubated for 15 min prior to blotting as described earlier. EcPKS2 data collection included additional super-resolution movies collected at 0.053 Å/px on the same microscope system as EcPKS1. The particles curated from this dataset were merged with the larger, but lower resolution, 1.06 Å/px dataset to yield a set of 264K particles (Supplementary Fig. 3). De novo volume creation and C1 NU-refinement in Cryosparc (47) resulted in a 3.15Å reconstruction of the full-length EcPKS2, where the bias of the map towards the larger modifying region left the dynamic condensing domain at lower nominal resolution (>6Å) (Supplementary Fig. 3 and 5). The model built into the full-length map includes the linker regions which was built in COOT (48), real-space refined and optimized. Running Cryosparc 3DFlex on this map using default settings, revealed conformational heterogeneity (Supplementary movie) (49). Masking, implementing C2 symmetry, a local and Global CTF refinement followed by NU-refinement produced the final condensing and modifying region maps at 3.03Å and 2.92Å resolution. Again, a ColabFold model was docked and rigid-body fit into the map followed by real space refinement in both COOT (48) and PHENIX (50) to generate the final model analyzed with MOLPROBITY (51) (Supplemental Table S1). Figures were generated with PYMOL (52) and CHIMERA (53).

##### 1.2.3 EcPKS2(AC) structure determination

Benzoyl-CoA (1 mM) and NADPH (1 mM) were incubated for 15 minutes with EcPKS2 at 6 mg $ml^{-1}$ in buffer with 0.1% Tween 20 and processed similar to other cryoEM grids. However, benzoyl-CoA was not found in the final model. Data was collected at 81,000x on KRIOS 4 at the

Pacific Northwest Center for CryoEM equipped with K3 camera, FFI and GIF for a final pixel size of 1.059Å/px. A total of 40 e-/$Å^2$ was recorded over 40 frames/exposure with defocus range of -1.0 to -2.0 and a slit of 10eV using SerialEM (54). After initial movie curation, 6,242 images were used to pick an initiate set of 614,919 particles which were reduced to 168,733 after 2D classification. The data were subjected to either a condensing or modifying region mask and further refined. Three-dimension classification of both masked sets identified differences in ACP occupancy at either the KS or DH domains and higher occupancy and classes with higher ACP occupancy were selected for final refinement (Supplemental Fig. 4. and Table S1).

### 1.3 Enzyme assays and kinetics

#### 1.3.1 General assay conditions

All assays used a buffer comprised of sodium phosphate (100 mM) and NaCl (100 mM) at pH 7.5. All enzymatic reactions were performed multiple times, and those reported herein used triplicate replicates.

#### 1.3.2 Quantitation of enzyme assay results

Reaction mixtures were centrifuged at 28,928 × *g* at for 10 min, and the supernatant (2 μL) was injected onto an Acquity UPLC protein BEH C4 1.7 μm column (2.1 x 100 mm) connected to a Water Xevo G2-XS Q-Tof. A linear gradient was employed from 5-100% mobile phase B (solvent A, $H_2O$ 0.05 % formic acid; solvent B, MeCN) at 0.4 ml/min over 6 min. MS method *a*: negative ion, sensitivity mode, MS range 100-1000 Da, scan every 0.1 s. MS method *b*: positive ion, sensitivity mode, MS range at 100-1000 Da, scan every 0.1s was used. Quantification was performed by comparing the AUC (area under curve) of the analyte with that for the internal standard, leucine enkephalin (55). Determination of kinetic parameters, $t_{1/2}$, and $IC_{50}$ was performed using Prism. To determine $k_{cat}$, one phase decay and log(inhibitor) vs normalized response model were used separately (56).

#### 1.3.3 PKS general characterization

PKS proteins (2.5 μM) were incubated with methylmalonyl-CoA (2 mM), NADPH (1 mM), and various acyl-CoA derivatives (1 mM) in 25 μL total volume, keeping reactions in dark at 22 °C for 18 h (Supplemental Fig. 7A).

**$^{13}$C-malonyl-CoA incorporation**

PKS proteins (2.5 μM) were incubated with methylmalonyl-CoA (2 mM), NADPH (1 mM), malonyl-CoA synthesis enzyme: MatB (5 μM) (57), $^{13}C_3$-malonic acid (1 mM), CoA (5 mM), ATP (7 mM), $MgCl_2$ (7 mM) in 25 μL volume, keeping reactions in dark at 22 °C for 18 h (Supplemental Fig. 9B).

#### 1.3.4 Time dependence of malonyl-CoA inhibition

PKS enzymes (2 μM) were incubated with NADPH (1 mM) at 22 °C for 1 min in 9 μL reaction volume; 1μL malonyl-CoA (1 mM, 0.5 mM, or 0.25 mM) was added, and the resulting mixture incubated for different times (1 min, 2 min, 4 min, 10 min, 20 min, 40 min), after which methymalonyl-CoA (1μL, 1 mM) was added. After 10 min, the reaction was quenched with acetonitrile (20 μL containing 10 ng/μL leucine enkephalin). The resulting mixture (10 μL) was further diluted with acetonitrile (40 μL) for analysis. Compounds **1** and **1'** were quantified to obtain the reported results (Supplemental Fig. 7).

#### 1.3.5 Determination of malonyl-CoA $IC_{50}$

PKS enzymes (9 μL, 2 μM) were incubated with different concentrations of malonyl-CoA (100 μM, 20 μM, 10 μM, 1 μM, 100 nM, 10 nM) for 40 min before adding methylmalonyl-CoA (1 μL, 1 mM). Reactions were quenched by 20 μL acetonitrile (with 10 ng/μL leucine enkephalin). The resulting mixture (10 μL) was further diluted with acetonitrile (40 μL) before analyzed by LC-MS using method *b*. Compounds **1** and **1'** were quantified to obtain the data shown in Fig. 5F.

#### 1.3.6 Kinetics experiments for EcPKS2 and mutants

PKS enzymes (10 μL, 2 μM) in sodium phosphate buffer were incubated with different concentrations of methylmalonyl-CoA at 22 °C for 10 min before quenching with acetonitrile (20 μL with 10 ng/μL leucine enkephalin). The resulting mixture (5 μL) was further diluted in acetonitrile (40 μL) with leucine enkephalin for LC-MS and analyzed by LC-MS by method *a*. Compounds **1, 3** and **4** were quantified to obtain data (Supplemental Fig. 5B).

#### 1.3.7 Kinetics experiments for EcPKS1 and mutants

PKS enzymes (10 μL, 3.6 μM) in sodium phosphate buffer were incubated with different concentrations of methylmalonyl-CoA at 22 °C for 30 min before quenching with acetonitrile (20 μL with 10 ng/μL leucine enkephalin). The resulting mixture (5 μL) was diluted in acetonitrile (40 μL) with leucine enkephalin and analyzed by LC-MS by method *a*. Compound **2** was quantified (Supplemental Fig. 5B).

#### 1.3.8 Structure determination of compounds

Most compounds described in this study were previously reported, and MS data here were directly compared with our previous study (17). In the case of internal malonate-incorporated compounds such as **6**, the reported structure is putative due to the low amount and instability of compounds. However, the incorporation of the number of malonate vs methylmalonate units was verified using $^{13}C$ labeling, providing high confidence for the number of each unit being incorporated but not their order in the resulting structure.

### 2 Additional descriptions of inactive domains of EcPKS2

#### $MT^0$ domain

A 14-residue linker, containing a beta-strand integral to the KR domain beta-sheet, connects the DH domain to an inactive methyltransferase-like ($MT^0$) structural domain, which sits at the periphery of the modifying region (Fig. 1). The remnant helical substrate-binding domain is followed by the pseudo-nucleotide-binding domain. In active methyltransferases the canonical *S*-adenosyl-methionine (AdoMet) binding residues E/DXGX<u>G</u>XG have evolved to EVGA<u>A</u>RG (EcPKS2) and EAGA<u>A</u>KG (EcPKS1) where the dihedral angle restrictions on the Ala residue flip the peptide chain to place the following Arg or Lys side chain in the AdoMet pocket (58). The $MT^0$ residues 1284-1300, 1338-1332, and 1458-1460 pack against the KR domain and contribute to its substrate-binding pocket, while 1321-1334 pack against the C-terminal linker between the KR and the ACP. The chain leaves the $MT^0$ domain and enters the $_{\psi}$KR structural domain prior to another

14-residue linker to the $ER^0$ domain. Overall, these observations are quite similar to those reported in the mFAS structure and support the close structural relationships between the AFPK and FAS enzyme families.

**$ER^0$ domain**

An inactive ER ($ER^0$, residues 1601-1958) domain packs against the initial DH domain and completes the dimer interface with DH:DH', $ER^0$:$ER^{0}$' and DH:$ER^{0}$' interactions. The domain arrangement and packing are homologous to mFAS and iPKSs such as LovB. Despite a mostly intact NAD-binding pocket, the EcPKS2 active site contains several mutations that would interfere with the binding of NADPH (Leu1658, Phe1856 and Arg1929), and the active site Lys and Asp have been mutated to Phe1856 and Phe1876 (Ser1842 and Leu1867 in EcPKS1). The occlusion of NADPH from the binding pocket suggests a possible reason for the lack of ER inactivity in the enzyme, and the resulting presence of a polyene in enzyme reaction products.

### Table S1: Structure statistics

| EM Databank Accession ID | EcPKS1-Condensing | EcPKS1-Modifying | EcPKS2(MC)-Condensing | EcPKS2(MC)-Modifying | EcPKS2(MC)-Full-length | EcPKS2(AC) Condensing | EcPKS2(AC) Modifying | EcPKS2(AC) Full-length |
|---|---|---|---|---|---|---|---|---|
| PDB Accession ID | | | | | | | | |
| **Data Collection/Processing** | | | | | | | | |
| Nominal Magnification | 81,000x | | 81,000x and 130,000x | | | 81,000 | | |
| Defocus Range (μm) | -0.08 - -1.6 | | -0.08 - -1.6 | | | -1.0 – 2.0 | | |
| Number of micrographs | 4,927 | | 17,420 + 2,838 | | | 6711 | | |
| Initial particles | 456,870 | | 965,000 | | | 614,919 | | |
| Symmetry imposed | C2 | | C2 | | C1 | C1 | | |
| Final Particles | 143,694 | | 258,974 | | | 106,739 | 93,476 | 168,733 |
| Map Resolution FSC 0.143 | | | | | | | | |
| Masked | 4.3Å | 3.5 Å | 3.0 Å | 2.9 Å | 3.2 Å | 3.0 | 3.1 | 3.1 / 6 Å |
| Unmasked | 6.4 Å | 4.2 Å | 3.7 Å | 3.4 Å | 3.6 Å | 4.0 | 3.4 | 3.24 Å |
| **Validation** | | | | | | | | |
| Map Correlation coefficient | 0.65 | 0.83 | 0.86 | 0.86 | 0.83 | 0.85 | 0.80 | 0.56 / 0.65 |
| Atoms | 12598 | 19482 | 14,588 | 20,890 | 34700 | 13234 | 43326 | 57300 |
| Protein residues | 1689 | 2512 | 1,870 | 2,602 | 4403 | 1724 | 2747 | 4571 |
| Bonds (RMSD) | | | | | | | | |
| Length (Å) | 0.014 | 0.010 | 0.008 | 0.010 | 0.009 | 0.005 | 0.005 | 0.006 |
| Angles (°) | 1.49 | 1.22 | 0.69 | 1.61 | 1.33 | 0.72 | 0.79 | 1.12 |
| MolProbity score | 2.59 | 2.39 | 2.39 | 2.87 | 2.81 | 2.12 | 2.22 | 2.27 |
| Clashscore | 31.51 | 17.47 | 30.11 | 19.25 | 23.62 | 14.12 | 17.03 | 22.93 |
| Ramachandran plot (%) | | | | | | | | |
| Outliers | 0.59 | 0.32 | 0.32 | 0.93 | 0.55 | 0.0 | 0.18 | 0.18 |
| Allowed | 9.33 | 11.98 | 6.23 | 7.00 | 6.86 | 4.26 | 7.73 | 6.18 |
| Favored | 90.08 | 87.70 | 93.45 | 92.07 | 92.59 | 95.74 | 92.09 | 93.64 |
| Rotamer outliers (%) | 1.21 | 1.15 | 0.89 | 6.24 | 4.32 | 1.67 | 0.68 | 0.70 |
| CaBLAM outliers (%) | 4.26 | 4.58 | 2.16 | 2.41 | 2.24 | 1.98 | 2.54 | 2.40 |

**Table S2: Plasmids used in this study.**

| Plasmid | enzyme | Refence |
|---|---|---|
| pxw55-EcPKS2 | EcPKS2 | [15] |
| pxw55-EcPKS1 | EcPKS1 | [1] |
| pEcPKSf2-1 | EcPKSf2-1 | [15] |
| pxw55-EcPKS2-A787V | EcPKS2- A787V | This study |
| pxw55-EcPKS2-A787H | EcPKS2-A787H | This study |
| pxw55-EcPKS2-Y706F | EcPKS2-Y706F | This study |
| pxw55-EcPKS2-S705A | EcPKS2-S705A | This study |
| pxw55-EcPKS2-S705A-Y706F | EcPKS2-S705A-Y706F | This study |
| pxw55-EcPKS2-S2198V | EcPKS2-S2198V | This study |
| pxw55-EcPKS2- SWQ/VYR | EcPKS2- SWQ/VYR | This study |
| pxw55-EcPKS2-EE/GG | EcPKS2- EE/GG | This study |
| pxw55-EcPKS2-1-2 | EcPKS2-1-2 | This study |
| pxw55-EcPKS1-VYR/SWQ | EcPKS1-WQSM | This study |
| pxw55-EcPKS1-polySW | EcPKS1-polySW | This study |

**Table S3: Primers used to generate mutants**

| | |
|---|---|
| pEcPKS2-A787V | catcgagatcgctccggtgggtctgctccagtccgtg<br>cacggactggagcagacccaccggagcgatctcgatg<br>ctatatcgtaataccatatggctcctcaggaacaagc<br>gtgatggtgatggtgatgc |
| pEcPKS2-A787H | catcgagatcgctccgcatggtctgctccagtccgtg<br>cacggactggagcagaccatgcggagcgatctcgatg<br>ctatatcgtaataccatatggctcctcaggaacaagc<br>gtgatggtgatggtgatgc |
| pEcPKS2-Y706F | gcaacaacatcagcttccactcatctttcatg<br>catgaaagatgagtggaagctgatgttgttgc<br>ctatatcgtaataccatatggctcctcaggaacaagc<br>gtgatggtgatggtgatgc |
| pEcPKS2-S705A | caacagcaacaacatcgcctaccactcatctttc<br>gaaagatgagtggtaggcgatgttgttgctgttg |

| | |
|---|---|
| | ctatatcgtaataccatatggctcctcaggaacaagc<br>gtgatggtgatggtgatgc |
| pEcPKS2-S705A-Y706F | caacagcaacaacatcgccttccactcatctttcatg<br>catgaaagatgagtggaaggcgatgttgttgctgttg<br>ctatatcgtaataccatatggctcctcaggaacaagc<br>gtgatggtgatggtgatgc |
| pEcPKS2-SWQ/VYR | agtggagaaggtcatgggcgtggccgc<br>gcggccacgcccatgaccttctccact<br>cagcatcaaggactaccgatacaccgtagcagac<br>gtctgctacggtgtatcggtagtccttgatgctg<br>ctatatcgtaataccatatggctcctcaggaacaagc<br>gtgatggtgatggtgatgc |
| pEcPKS2-S2198V | agtggagaaggtcatgggcgtggccgc<br>gcggccacgcccatgaccttctccact<br>ctatatcgtaataccatatggctcctcaggaacaagc<br>gtgatggtgatggtgatgc |
| pEcPKS2-EE/GG | ctggtccaccatagagccgccccctgcggccacg<br>cgtggccgcagggggcggctctatggtggaccag<br>ctatatcgtaataccatatggctcctcaggaacaagc<br>gtgatggtgatggtgatgc |
| EcPKS2-H903A | ctatatcgtaataccatatggctcctcaggaacaagc<br>actacttgttggaggctgcagtggatgggcgcatgt<br>acatgcgcccatccactgcagcctccaacaagtagt<br>gtgatggtgatggtgatgc |
| EcPKS2-L1075S | caggcactcaccaggccagtcccaccctgctcga<br>gactcgagcagggtgggactggcctggtgagtgcctg<br>ctatatcgtaataccatatggctcctcaggaacaagc<br>gtgatggtgatggtgatgc |
| pEcPKS2-1-2 | ctatatcgtaataccatatggctcctcaggaacaagc<br>gacgagtccgagcccgaggccgaaccgtactcaaactc<br>cgaagagtttgagtacggttcggcctcgggctcggactc<br>cacgcccatagacttctccaccaggacgtggcaggcgac<br>ggttgtcgcctgccacgtcctggtggagaagtctatgggcgtg<br>gtgatggtgatggtgatgc |
| pEcPKS1-VYR/SWQ | ctctggcaaggactggcagtacaccgtgggtgac<br>gtcacccacggtgtactgccagtccttgccagag<br>gtgatggtgatggtgatgc<br>ctattaactatatcgtaataccatatggcgcccaataacacccagcaag<br>ctccacagcggccatgctcttctcggccag<br>ctggccgagaagagcatggccgctgtggag |

| | |
|---|---|
| pEcPKS1-polySW | ctcgccctccacagcgctgctgctgctctcggccagga<br>cctggccgagagcagcagcagcgctgtggagggcgag<br>ccacggtccaccaccaccacttgccagagaacttggc<br>ctctggcaagtggtggtggtggaccgtgggtgacgct<br>ctccacagcggccatgctcttctcggccag<br>ctggccgagaagagcatggccgctgtggag |

**Table S4: Sequence of EcPKS2-EcPKS1-EcPKS2 hybrid enzyme**

| **Sequence (EcPKS1; EcPKS2; EcPKS2 ACP linker)** |
|---|
| **MAPQEQASSSQQDDAPPKTPNVVEPYKGEVAICGLSGRYPESANVGELEYNLFNK<br>IDMVTIDNRRWEPGYLGTPERMGKVKTITDFDAEFFGVHTKGAQTMDPMLRNL<br>LEVVYEAIVDAGESLESMKGTRTGVYIGVSNNEVDTAYMKNWTDDDAYMVQGC<br>HHSMYPNWISFFFDFSGPSTAYNTACSTSLVCLDAAERHLRMGVIDNAIVGGSNFI<br>YRPATTKLFMGMNFLGSSTCKAFDESGDGFVRGEVASAILLKKADTAKRVYCTLV<br>GSMLNNDGNQTNGILYPNSEAQEQLMTDIYSTHKIDANEVKYFECHGTGTQAGD<br>PNETRAICNAVCKGKKDPLLIGSIKSNLGHGETASGINGISKVIITMHSRQIPPNLH<br>FKNPNPKIPGLFDGRLKVVTETTPFDGGLIAINSFGMGGTNAHAIFRSFDKRAEPH<br>PASDKPRLFTYCARTEEGLQKIFEEAHKHASNVEFHALCQESANTKPKSLPYRGA<br>TILNAEGEYTEIQKCPSKAREVWFVYSGMGSQWVGMGRSLMALDVFRQSIEETA<br>AILSPFGVDLMSLLMDGTEDKLKEIMPPFICINAIQLALTDLLNSMGIVPDGLVGH<br>SLGEVGCAYADGCLTRREAILSAFWRAKAVIDCEVKPGKMAAVELTWEEAKRLC<br>PPGVVAACHNSQDSVTISGGAQEMTKFMAELSAQGVTVKEVNSNNISYHSSFMTE<br>PAAYLKKGLEKEIVPKPRSKKWISTSIPEERWGNPEAQTADASYQANNLLSSVLFY<br>EGLQKIPSNAIAIEIAPAGLLQSVIKKSLGQDCTIVALQKRKSPNNLEVFFSALGKC<br>YSHGVPMNPLGLYPAVQFPVSIDTPMLSSMVSEAWDHSAKWRVPLVEEFEYGSAS<br>GSDSSYSIDVSADSPDRYLLDHQVDGRELFPACGCLVLAWKTLAALNGRDFEQMP<br>VRLSRVEIHQAMFLPKSGSATVTVSVMPRTGEFQVCENENLLASGFVTCPDKDVL<br>ETSTHAQTRSSLQDRPATEVLTRDEVYRELILRGYEYGPYFQGILRASVDGQESEI<br>TWDGRWVSFMDSVLQMDILARPGDYQMLPIKFQSINIDPRVQPAAPAEDEDVVV<br>LPGRFDPVLDIVSAGGVEIRGLETISASRRLTHAPEVVEEYRFVPHHVTGRDDPGA<br>KRPGATVDIREYADACLAFAVQGIKKWLSEDKDKVLPQKDLLQDALGLANQDLG<br>SKSSSSDFISAKAALERILKQQNGHQQHGFGLFHTLNLAFSEPLEIGFRETLKNKI<br>HHMRYDMWDDCLMSAVECADSLKLCIDTVAENTTSHIVNVLEAGAAKGAFYRR<br>AIPEALAKFSGKDYRYTVGDASPMDDAKEFSVKTLQFDAYDPANFPASQAHAHDL<br>LVLKWVLHQQEDLDAAMAGFCGFVRPGGFILVQEFVHRLPTLLAVEAVTDHPLP<br>RKSGDRVLGRYYSAAQWRELFRRHGLVEVIHRSDGALADMFLLRSRVEVMTPPT<br>VLHLDDLSCSWLEEVKAKYSDLEAMPQDARLWLVGKSDCNGMLGFFNCLRQEP<br>GSERVRCVQVCGDSVPDLSPGSAEFKYLAEMDLAFNVHKDGKWGVYRHLAITD** |

**DQRRQQFPTEHAFVDTLTSGDLSTLTWVRSPLNLHASSEKGQDCELCTVYMAGV**
**VSRDLALACGKLRRDELPAGMFCKEGTLGIEFSGRDTKGKRVMGLCAPPALASS**
**VLCLRSSLWSVPQHWSLEEAATVPVAYSTAYYALVIRGHVRPGDTVLVHAGGSPV**
**GQAAIAVAQSCGCEIFISTATDAETSSLKSMFPRLKDRNFCSCKDASFERHVKKET**
**SGKGVDIILNCTTGELLGASIRLLASRGRFLNLASGRGSDAELVFSGSGRRDTSFH**
**DINLDTLIDAQGPEWTELTSLVQKGIQSGLVKPLARTVYAMDRLVDVFKLLEEGA**
**QAGKLLVKIREEEAEKITLPAKKTFEAVPRTFFHPAKSYVIVGGLGGFGLELAHW**
**MVLRGVRKLVLTSRNGITTGYQTRKIAFLRSLGADIVVCAVNVTSQAAADRLVKT**
**ATDLGPLGGVFNLGLNLRDALLVEQTAENYKQTLEAKIQTTSLLDGISRSPKIQPT**
**LDHFVMFSSLSAGHGIPGQTNYGWGNSYMDRLCEKRRAQGLPGLSIQWASIADV**
**GFVGTKGNNVVIEGKWPQRMYNCLQVCDYFLSQNRPVVACHVLVEKSMGVAAG**
**EESMVDQVLRAVGKVLGIKDVSSVDGDKEFIDMGVDSLMSVEIKQALERDAGLVI**
**STKDTQLMTFNTLRSMVKGS**

### Supplementary Figures

**Supp. Fig. 1. Schematic of full catalytic cycle for EcPKS2 (in green) and EcPKS1 (in orange).**

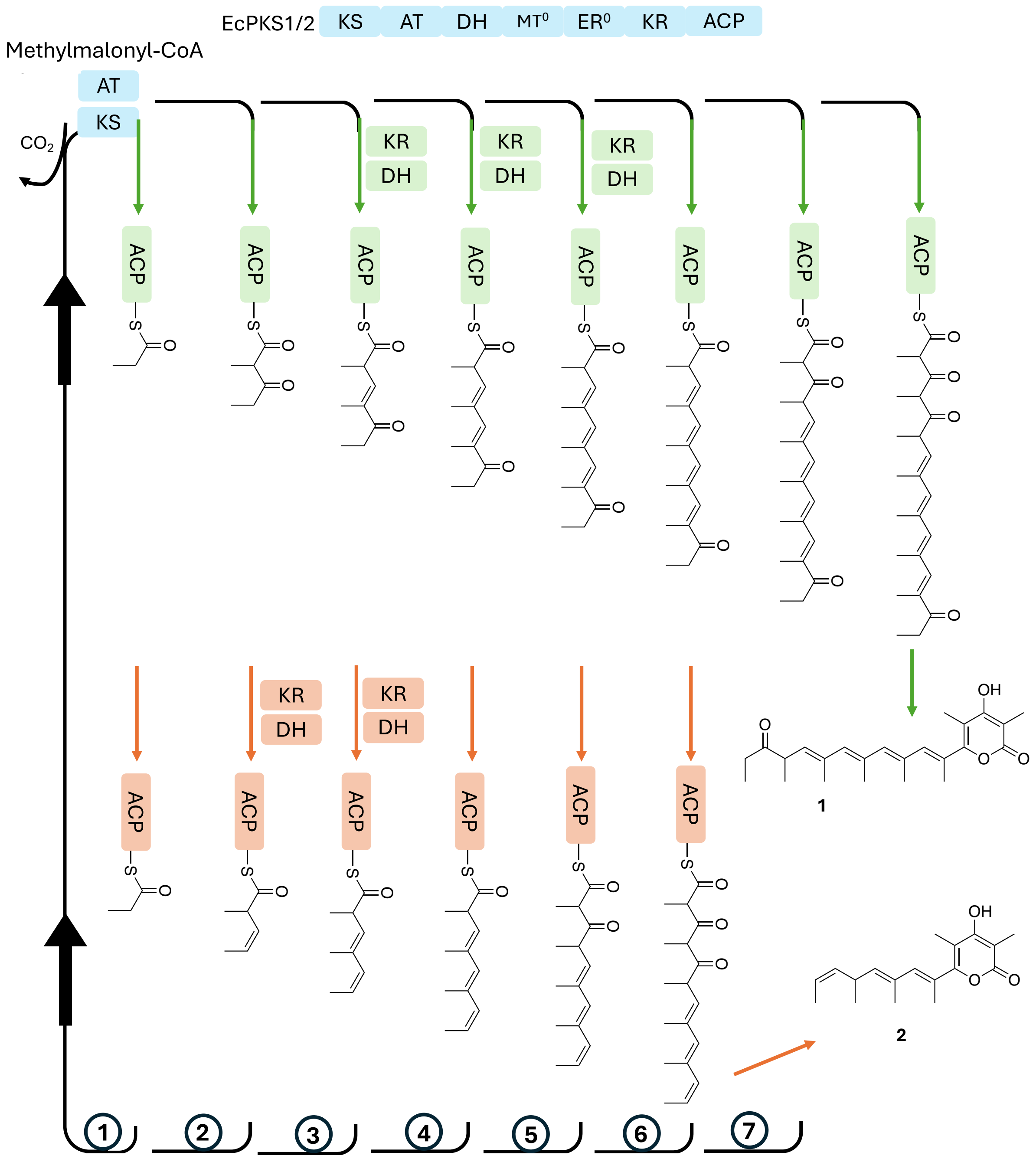

**Supp. Fig. 2. Structure determination method schematic for EcPKS1.**

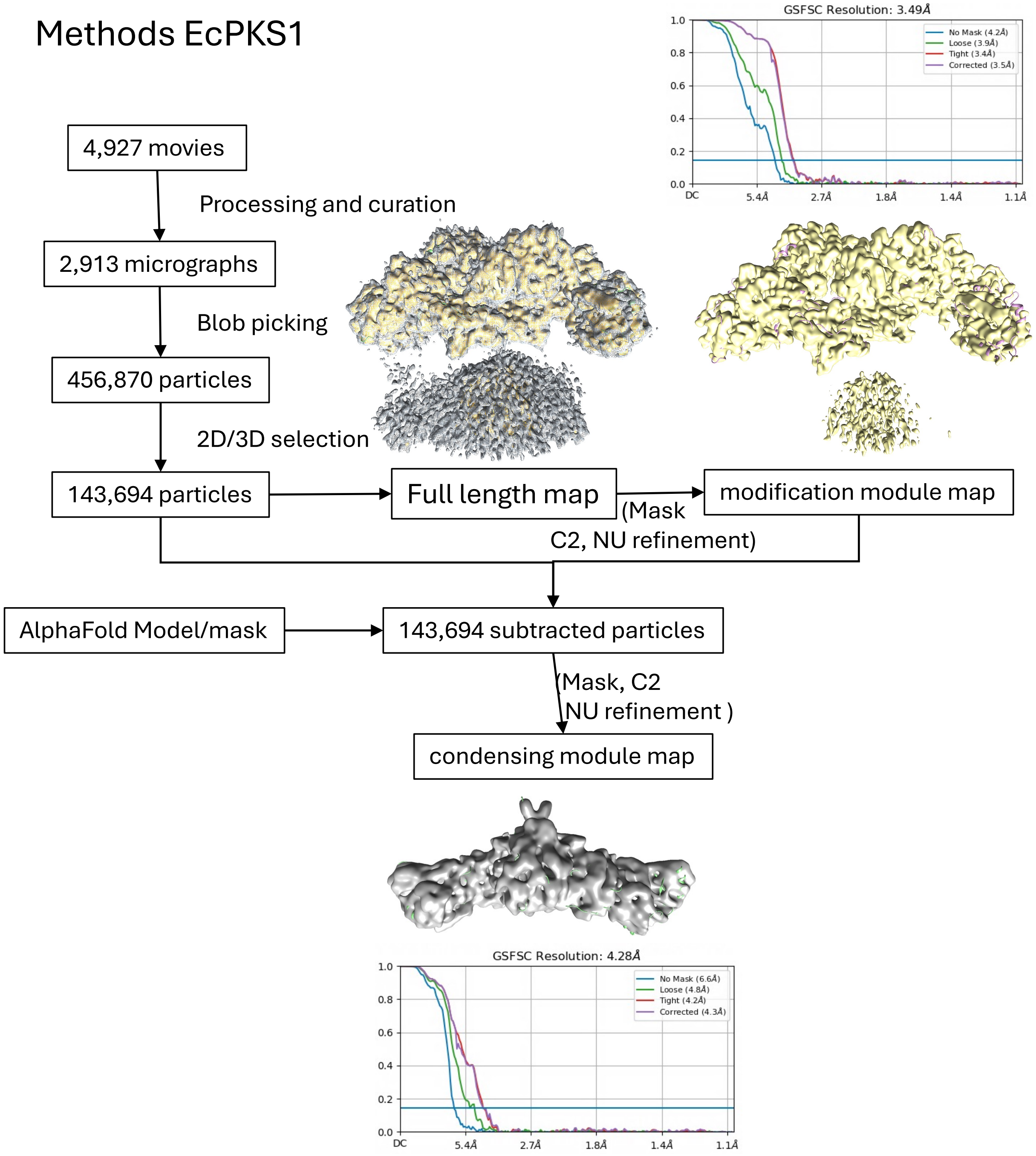

**Supp. Fig. 3. Structure determination method schematic for EcPKS2(MC).**

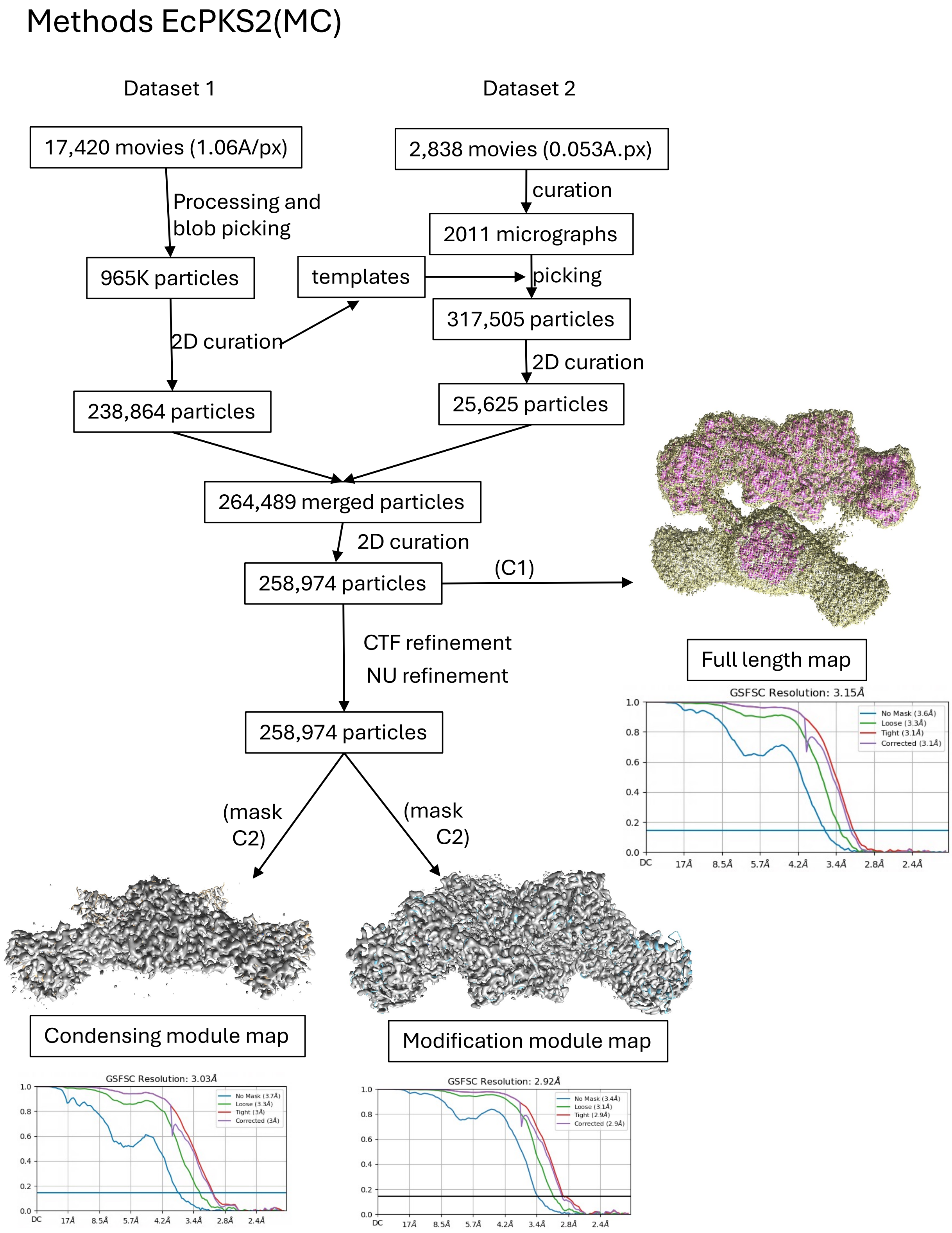

**Supp. Fig. 4. Structure determination method schematic for EcPKS2(AC).**

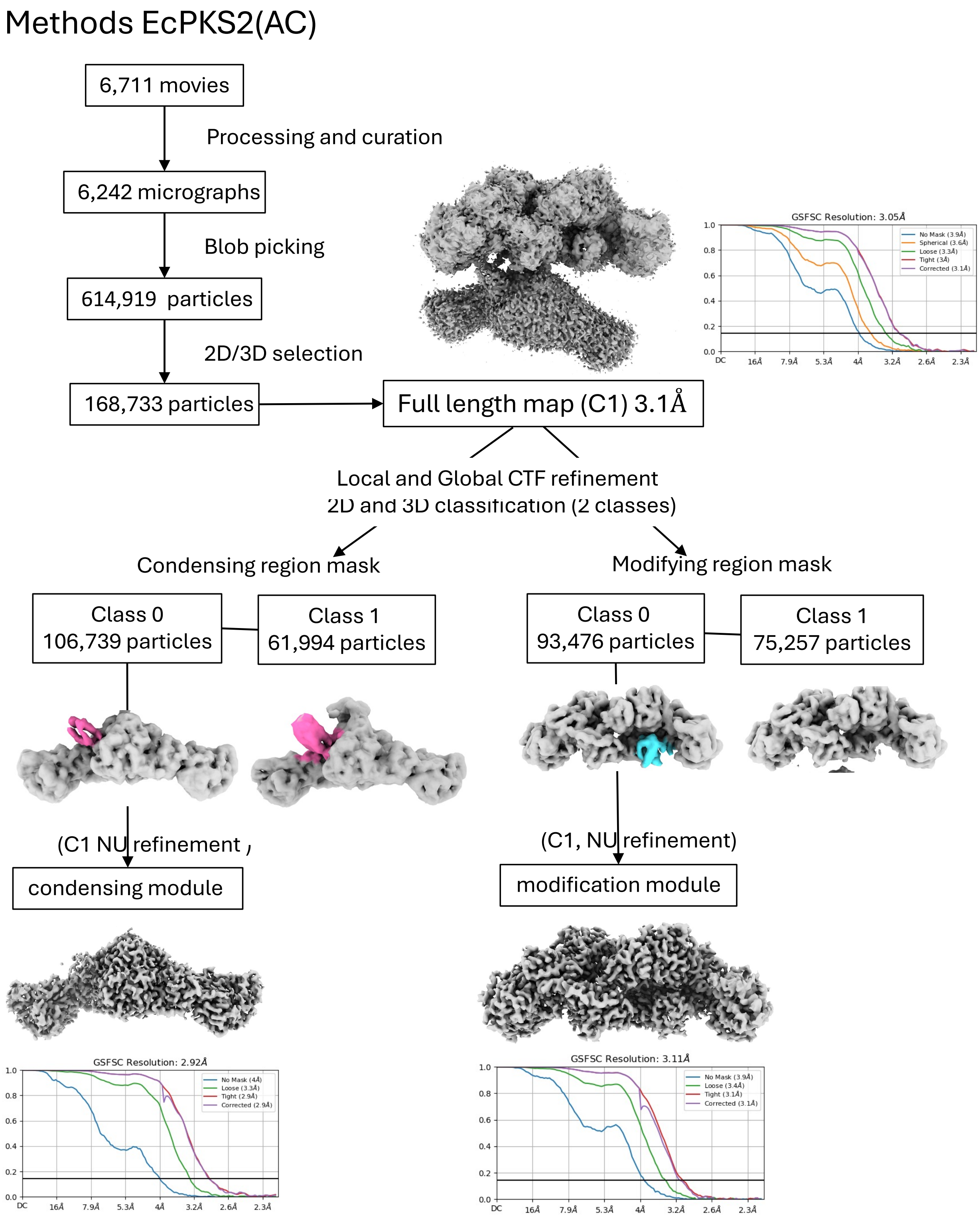

**Supp. Fig. 5. Local resolution maps for EcPKS1, EcPKS2(MC) and EcPKS2(AC) colored from high resolution <3.0Å (cyan) to low resolution >6Å (white). A)** Diagram of the domain arrangements. **B)** Independently refined EcPKS1$^{C2}$ condensing (bottom) and modifying (top) regions. **C)** Full-length EcPKS2(MC)$^{C1}$ refined map. **D)** Independently refined EcPKS2(MC)$^{C2}$ condensing (bottom) and modifying (top) regions. **E)** Full-length EcPKS2(AC)$^{C1}$ map and **F)** independently refined EcPKS2(AC)$^{C1}$ condensing (bottom) and modifying (top) regions.

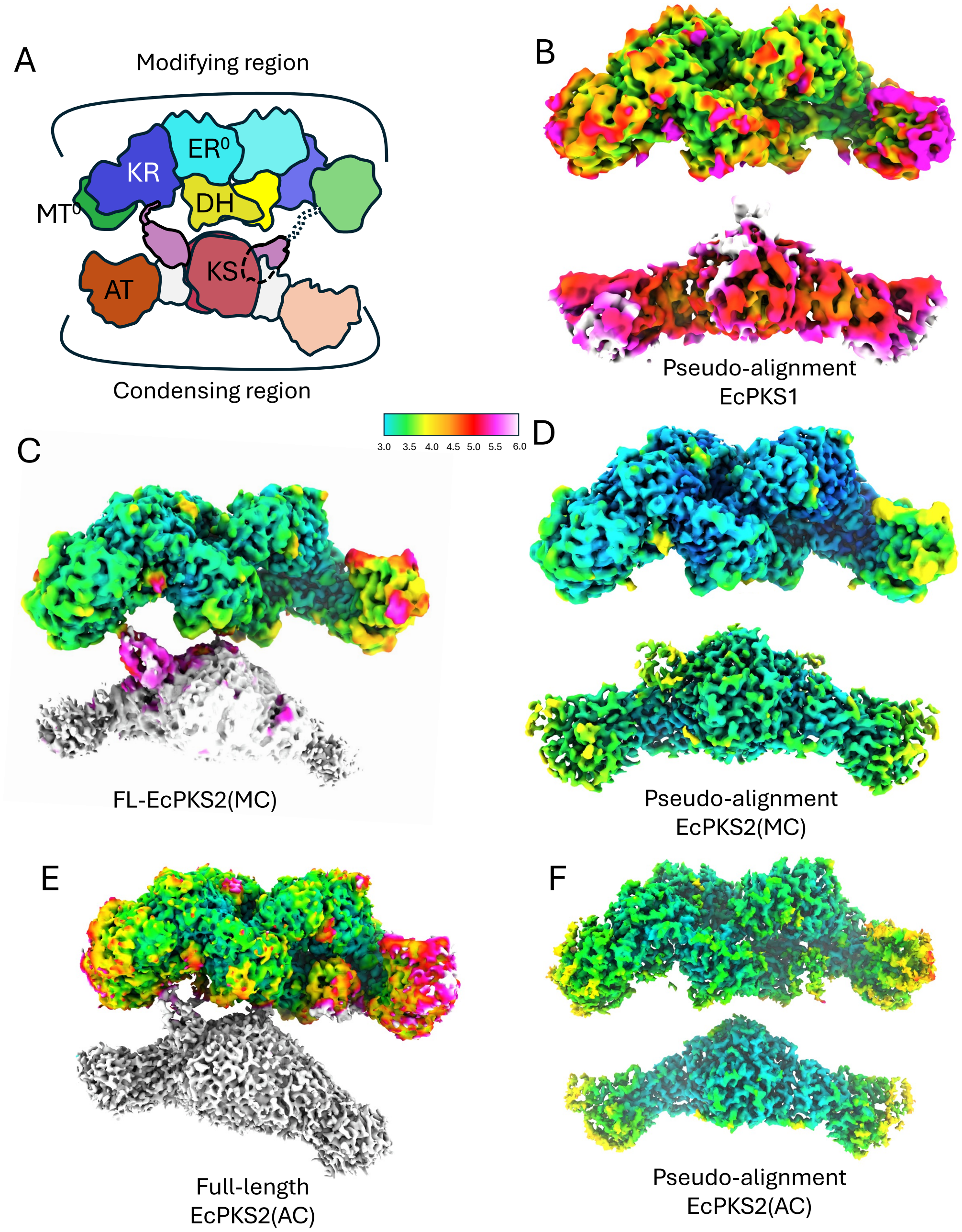

**Supp. Fig. 6. Representative density. A)** MC bound to the EcPKS2(MC):AT domain (orange). **B)** ACP-pPant (purple/white) docked at the EcPKS1(MC):KS domain (red). **C)** Acetyl-Cys a the EcPKS2(AC):KS active site (red). **D)** NADPH in EcPKS2(AC):KR domain (blue). **E)** ACP-acetyl-pPant (purple/white) docked at the EcPKS2(AC):DH domain (yellow). **F)** Full-length EcPKS2(MC) map at the ACP linker (pink) region, KR (blue), MT (green), ACP (purple).

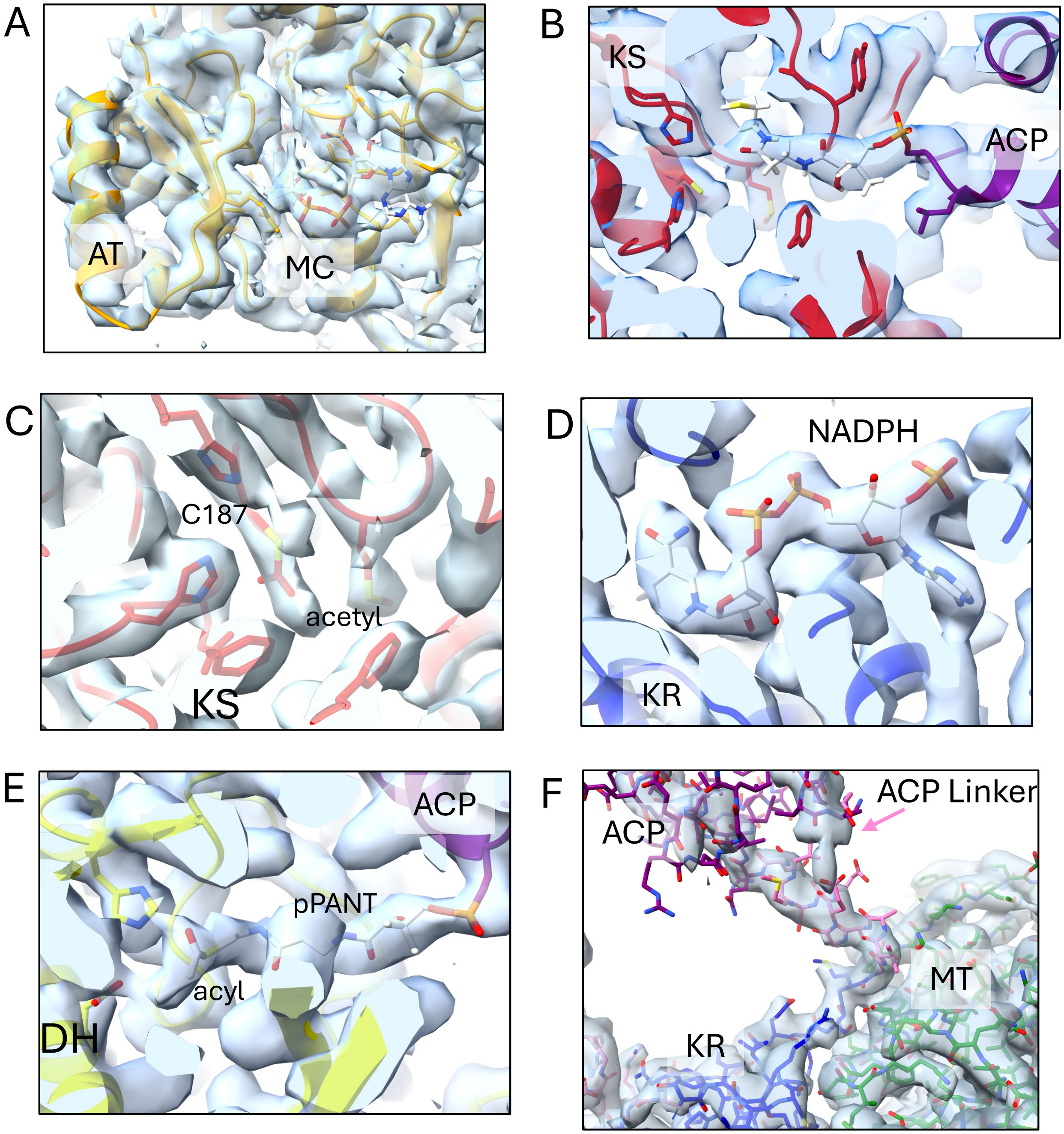

**Supp. Fig. 7**. **ACP linker function characterization. A)** Malonyl-CoA (MC) inhibition ($t_{1/2}$) of linker mutants with different concentrations of MC: 250 µM (green), 500 µM (blue), and 1000 µM (red). AUC represents compounds **1** and **1'**. Panels A1, A2, and A3 represent wild-type and two mutants, respectively. **B)** Catalytic efficiency decreases in ACP linker mutants meant to disrupt the MT-linker interface in EcPKS2, while putatively establishing that interface in EcPKS1 increases efficiency. **C)** Structures of products **1'** and **2'**, in which acetate derived from MC has been incorporated into the starter unit.

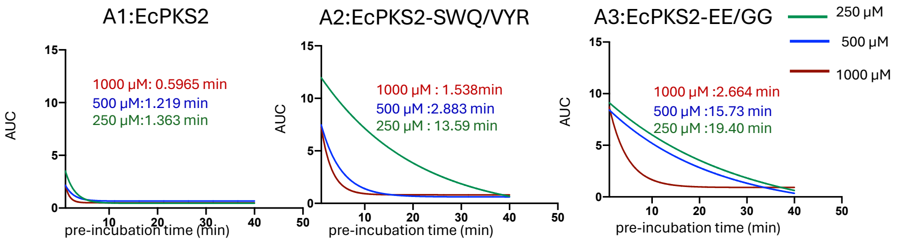

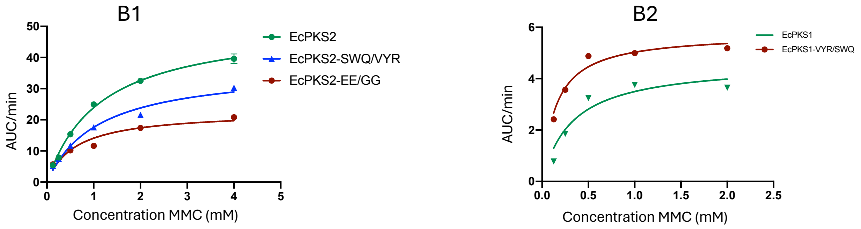

| | $k_{cat}$(min$^{-1}$) | $K_m$ (mM) | $k_{cat}/K_m$ (mMmin$^{-1}$) | $V_{max}$(AUCmin$^{-1}$) |
|---|---|---|---|---|
| EcPKS2 | 25.61±0.596 | 1.14±0.066 | 22.46 | 51.22 |
| EcPKS2-SWQ/VYR | 18.41±0.778 | 1.09±0.117 | 16.89 | 36.82 |
| EcPKS2-EE/GG | 11.37±0.598 | 0.61±0.097 | 18.64 | 22.73 |
| EcPKS1 | 1.285±0.106 | 0.323±0.084 | 3.98 | 4.626 |
| EcPKS1-VYR/SWQ | 1.600±0.139 | 0.146±0.056 | 10.05 | 5.759 |

C

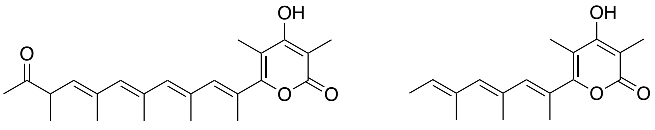

**1'** **2'**

**Supp. Fig. 8. ACP interactions. A)** Overlay of three ACP-docked AT structures with EcPKS1(MC):AT. VinK/VinL (PDBcode:5czd)[30] in yellow, Lsd14(apo-ACP) (PDBcode:7s6b)[43] in purple, SalAT9M-ACP9 (PDBcode:7vrs)[23] in green. **B)** Overlay of EcPKS2(MC):ACP-pPant with KS in red and ACP in purple, with Lsd14(ACP-pPant)(PDBcode:7s6c)[43] in green, GfsA(ACP-pPant)(PDBcode:8in9)[24] in yellow, and the yFAS(double ACP)(PDBcode:8psl)[33] in blue. **C/D)** Surface representation of the condensing region model (grey) with residues at the predicted AT:ACP interface highlighted in orange/magenta respectively. ACP in white. Rotation of 90 degrees along the x-axis between the two panels. **E/F)** Same as C/D but for interface with KS domain as modeled in EcPKS2(MC). **G/H)** Same as C/D but for interface with DH domain as modeled in EcPKS2(AC).

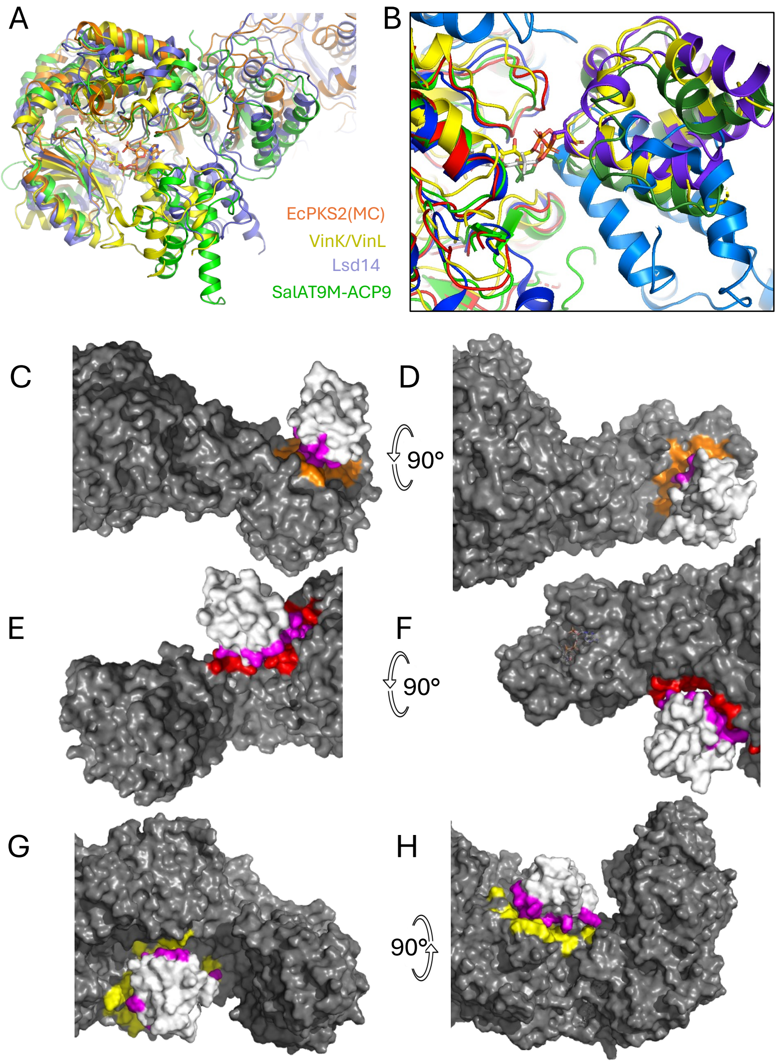

**Supp. Fig. 9. Domain fusion experiments further verify function of ACP linker. A)** EcPKS2-1-2 hybrid enzyme, designed with the new linker model, synthesizes unreduced product (**1**) in addition to **3** and **4**, while we reported previously that hybrid EcPKSf2-1 does not make **1**[17]. **B)** EcPKS2-1-2 hybrid enzyme incorporated MC as both starter and extender units to synthesize compound **6**; the corresponding structure and mass spectra are shown at right. In the spectra, when $[^{13}C]_3$-malonyl-CoA was used, an ion was observed at *m/z* 251.1395, while absent with unlabeled MC the observed ion was at *m/z* 245.1190. This result confirms that **6** contains 3 units derived from MC; their putative positions are shown in the bold bonds represented in **6**.

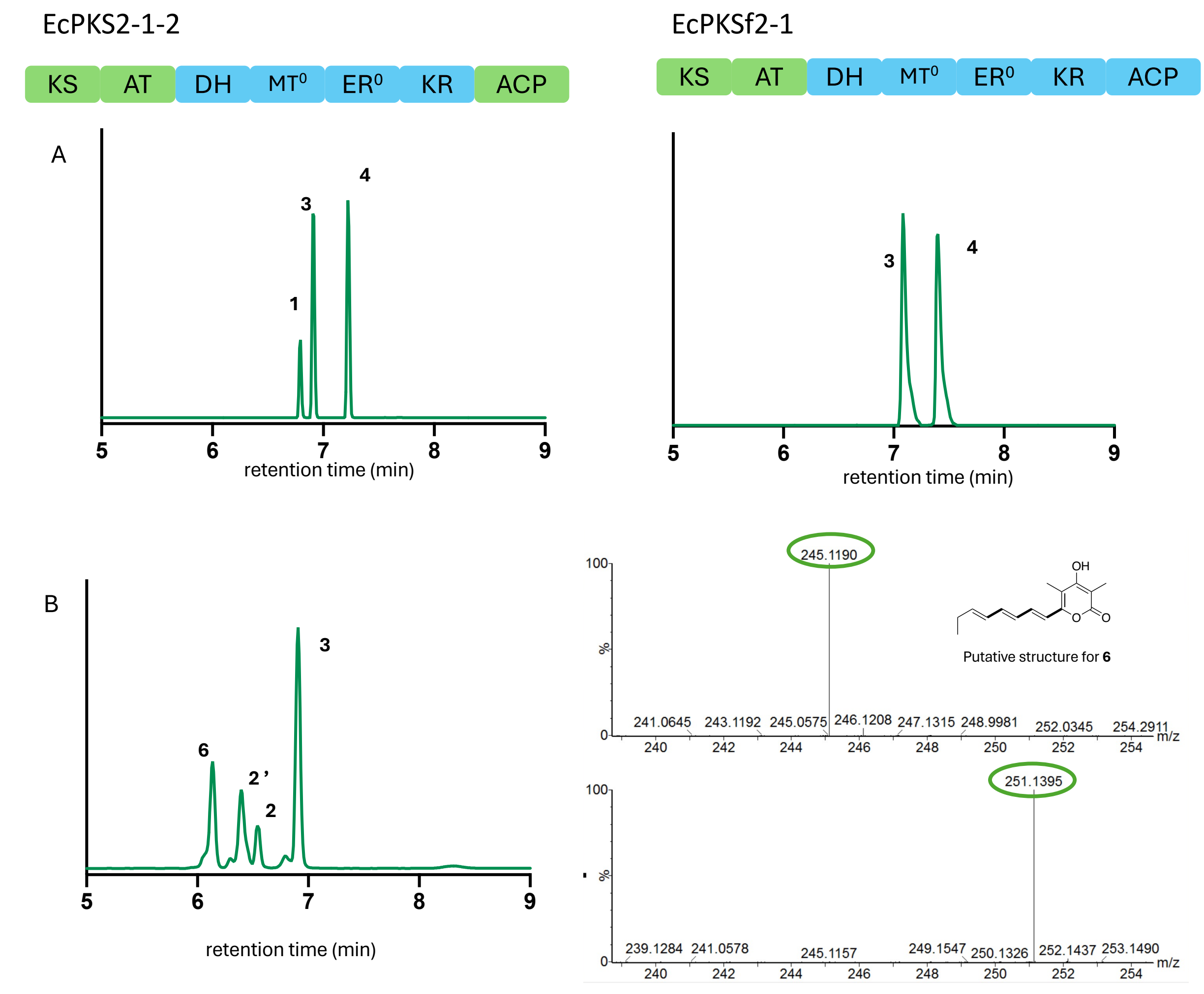

**Supp. Fig. 10**. **Characterization of EcPKS2 and EcPKS2 mutants. A)** The major products of EcPKS2 do not change when the AT is mutated or when the ACP linker is modified. **B)** When the EcPKS2 DH active site is disrupted, **1** is no longer produced, and instead short pyrones are synthesized, consistent with a lack of activity in the modifying domains.

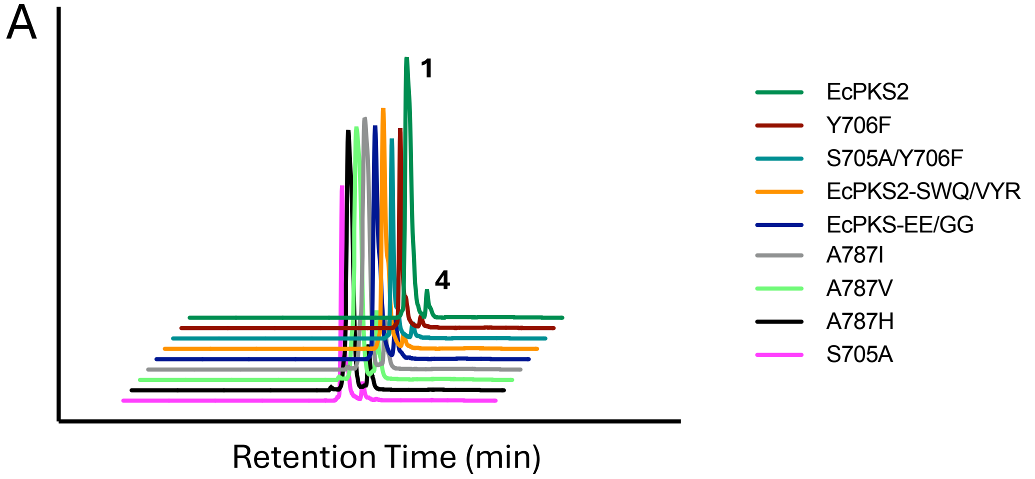

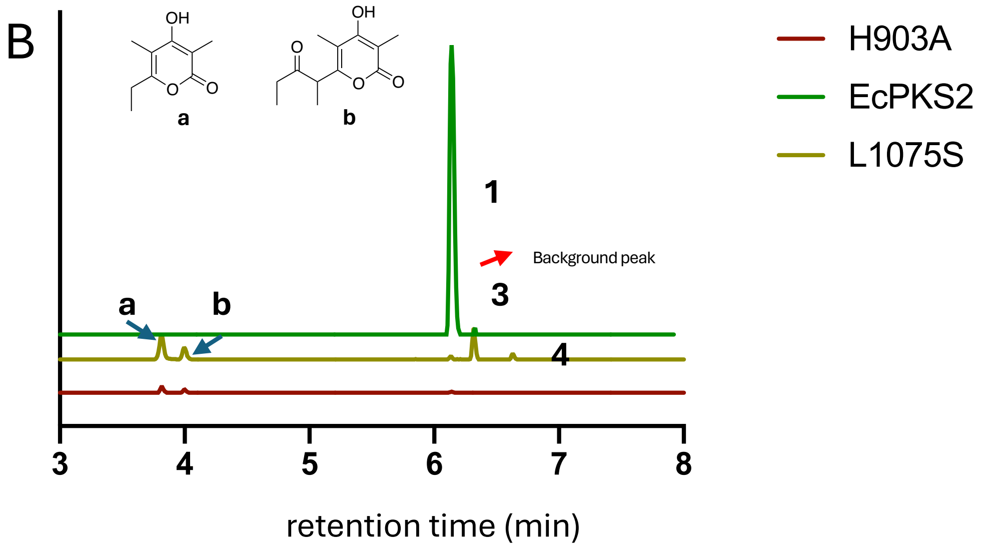
